# RUNX1 and YY1 modulate neuronal fate and energy metabolism in Alzheimer’s disease

**DOI:** 10.64898/2026.04.13.716801

**Authors:** Raffaella Lucciola, Joseph R Herdy, Yatavee Vajaphattana, Lukas Karbacher, Thais S Sabedot, Michael S Cuoco, Larissa Traxler, Austin Kang, Mack Reynolds, Jeffrey R Jones, Simon T Schafer, Jerome Mertens, Fred H Gage

## Abstract

Loss of neuronal identity and metabolic dysfunction are features of Alzheimer’s disease (AD), yet the upstream-acting molecular drivers remain incompletely understood. By integrating multi-omics data from patient-derived induced neurons (iNs) and AD post-mortem human brains, we discovered that AD neurons express two master transcription factors (TFs), RUNX1 and YY1. While these TFs are primarily expressed during development where they play fundamental roles in cell fate determination and cellular bioenergetics, respectively, they can be reactivated in adult neurons in response to stress. To understand their functional role in AD neurons, we overexpressed RUNX1 or YY1 in aged iNs and found that the expression of each TF was sufficient to recapitulate two AD-associated features. Specifically, RUNX1 overexpression caused loss of neuronal fate, whereas YY1 overexpression regulated gene regulatory programs associated with metabolic dysfunction. Conversely, downregulation of either TF, in AD iNs, reinstated gene regulatory programs associated with a healthy mature neuronal phenotype. Together, these findings identify two transcriptional master regulators of the AD neuronal phenotype and establish a mechanistic foundation for further studying their role in the pathogenesis of AD and as putative therapeutical targets for the treatment of AD and age-associated neurodegeneration.

## Introduction

Alzheimer’s disease (AD) neurons simultaneously lose mature identity and undergo maladaptive metabolic dysfunction, but the upstream transcriptional drivers linking these processes to aging remain poorly understood. Progressive biological aging appears to erode the neuronal epigenetic landscape, creating a permissive state in aged neurons in which disease-specific transcriptional programs can drive vulnerability and pathogenesis towards AD ^1–8^. Yet, the study of human age-associated diseases like AD at the cellular and molecular levels is challenging, due to the complexity of the brain and the difficulty of obtaining live human brain cells for functional studies. Findings from post-mortem human brains have accelerated our knowledge and understanding of the complexity of AD. Genomic, epigenomic and transcriptomic studies have shown large changes in the expression of genes specifying neuronal identity, including synaptic genes, and of genes involved in energy metabolism as well as early developmental and de-differentiation regulatory factors, whose binding site accessibility is altered, at least in part, by an age-dependent epigenetic drift ^1,2,8–16^. However, although these post-mortem human brain studies have mapped extensive changes in neuronal identity, developmental regulators, and metabolic states, they have not disentangled cause from consequence or directly tested how candidate regulators alter neuronal phenotypes.

The advent of stem cell research and cell reprogramming technology opened new venues and offered unprecedented research opportunities. Using direct cell reprogramming, we previously developed a novel approach to generate an age-equivalent neuronal model by transdifferentiating dermal fibroblasts obtained from patients into induced neurons (iNs) ^3,17^. Unlike induced pluripotent stem cell (iPSC)–derived neurons, iNs retain the aging signature of their donor and allow for modeling of human age and age-related diseases such as AD *in vitro* ^17–22^. We generated iNs from AD and age-matched control subjects and reported that AD patient-derived iNs recapitulate specific AD signatures observed in AD brains, thereby providing a valid system to model human AD-associated neuronal changes *in vitro* ^3^. In line with findings from *in vivo* studies, we identified specific gene expression and epigenetic changes that were associated with the induction of cellular phenotypes including mature neuronal identity, de-differentiation and metabolic shifts ^3,23^.

To advance our understanding beyond descriptive cataloging of dysregulated genes and genomic features, we have integrated multi-omics datasets from AD iNs and AD brains to identify age-associated transcription factors (TFs) that act upstream of AD pathogenetic changes. Our study highlights two early developmental and cancer-associated TFs, RUNX1 and YY1 ^24–30^, whose expression increases specifically in aged AD neurons *in vivo* and *in vitro*. Of note, RUNX1 is a TF that acts as a master regulator of cell fate determination and plays a key role in neuronal lineage specification during development ^26,31–38^. While expressed at low levels in the adult brain, RUNX1 can be reactivated following traumatic injury and during regenerative responses ^28,39^. On the other hand, YY1 is a ubiquitously expressed TF with fundamental roles in controlling cellular metabolic state during brain development ^40^. Similarly to RUNX1, YY1 can also be re-engaged in adult cells in response to damage or under pathological conditions and previous evidence has indicated that YY1 can drive metabolic reprogramming analogous to that observed in AD neurons. For example, in satellite muscle stem cells, YY1 drives an injury-induced shift towards glycolytic metabolism ^24^, whereas in cancer cells it promotes Warburg-like metabolic reprogramming ^41^, suggesting a conserved role in coordinating stress-induced bioenergetic adaptation. Previous studies from post-mortem AD human brain tissue have focused on RUNX1 and YY1 and suggested that they could have active regulatory roles in AD-related pathological dysfunction in neurons and microglia ^8,42^. Here, we hypothesized that AD neurons reactivate RUNX1 and YY1 as an adaptive response to AD-induced stress, and we have leveraged the iN model to examine how dysregulation of these TFs in AD neurons influences cellular phenotype.

## Results

### Identification of dysregulated TFs in AD neurons

AD neurons exhibit specific gene expression and epigenetic changes that are associated with the induction of cellular phenotypes such as fading neuronal identity, de-differentiation and metabolic shifts, among others. However, the drivers acting upstream of these changes remain incompletely understood. To address this question, we cross-referenced RNA-sequencing (RNA-seq) data from AD patient-derived iNs that we generated from a well-characterized cohort of AD patients and age- and sex-matched, non-demented control donors (*n* = 13 and *n* = 15, respectively) ^3^, and single-cell RNA-seq (scRNAseq) data from post-mortem AD brains, obtained from AD patients and cognitively non-demented controls from the Religious Order Study (ROS) and the Rush Memory and Aging Project (MAP) ^2^ (Figure 1A, left panel (i)). For this analysis, we specifically examined excitatory neurons from the AD brains, as they represent the cell type that iNs represent^3^.

**Figure 1.**
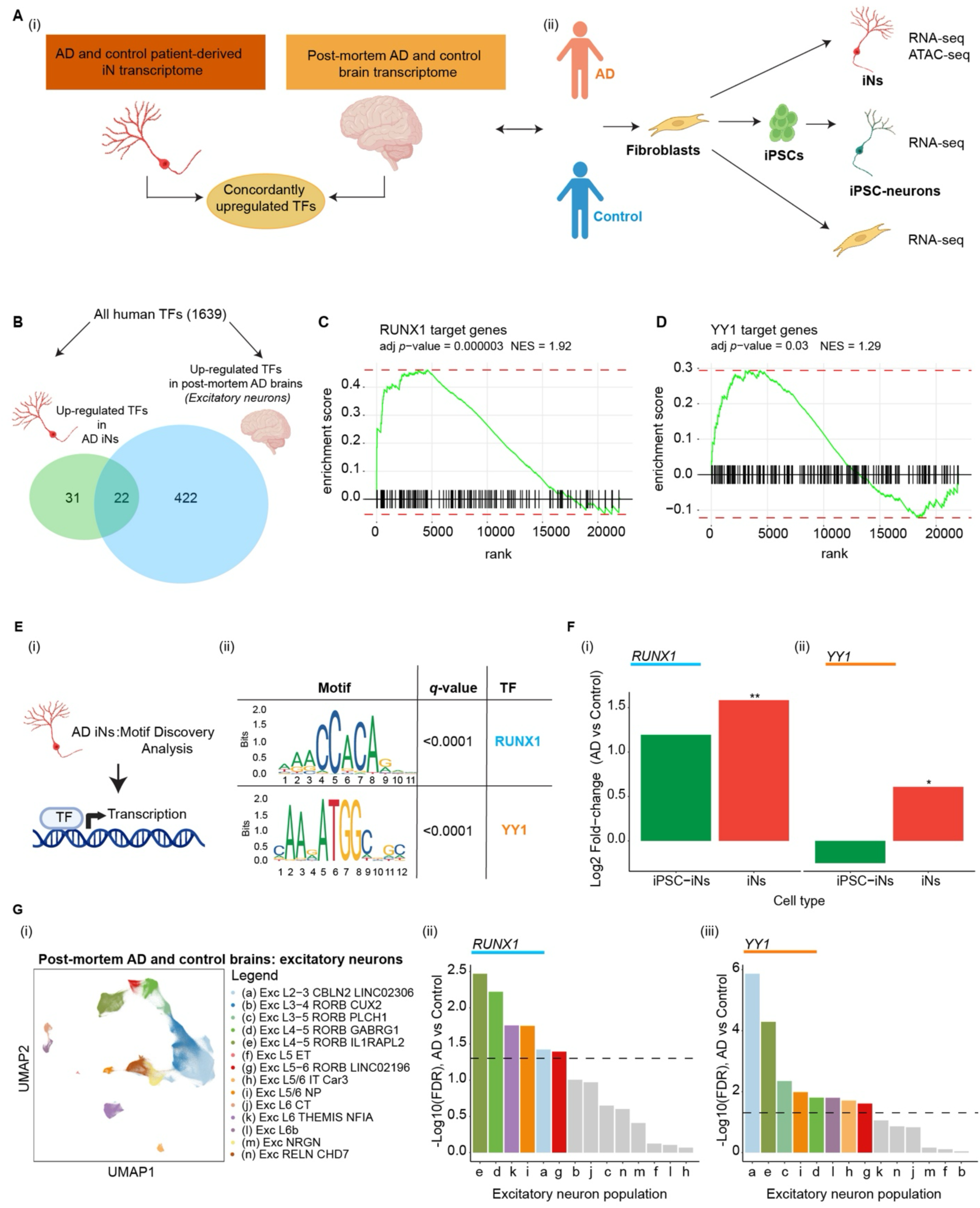
Identification of dysregulated TFs in neurons from AD patients. (A) Schematics. (i) Cross-referencing transcriptomes of AD and control iNs with human post-mortem AD brain transcriptomes; (ii) integrating these findings with complementary multi-omic datasets generated from AD and control iNs, iPSC-iNs and fibroblasts. (B) Venn diagram showing number of TFs significantly upregulated in AD iNs and excitatory neurons from human post-mortem AD brains (C-D) Gene set enrichment analysis (GSEA) of RUNX1 and YY1 target genes in excitatory neurons from human post-mortem AD brains. (E) HOMER motif discovery analysis of AD iN differentially open peaks captured significant enrichment of RUNX1 and YY1 binding motifs in AD iNs as previously published ^3^. (F) RUNX1 (i) and YY1 (ii) differential expression in AD iPSC-derived neurons (iPSC-iNs) and iNs. Adjusted *p* (*p*adj) values were calculated for differential gene expression using DESeq2 (* *p*adj < 0.05, ** *p*adj < 0.01, *** *p*adj < 0.001). (G) (i) UMA plots of excitatory neuron population as previously described ^2^ (ii-iii) Significance values of RUNX1 and YY1 across the excitatory neuron population from human post-mortem AD brains.

Cellular phenotypes are defined upstream by transcriptional regulators that reprogram the transcriptional and epigenomic profiles defining cellular fate and state. Therefore, we specifically searched for TFs. First, we extracted all human TFs that were significantly upregulated in AD iNs and in AD excitatory neurons and then searched for those that were concordantly upregulated across both systems. This analysis captured a set of twenty-two TFs (Figure 1B, Supplementary Figure 1A). We then examined the expression of their target genes as an indicator of TF activity and prioritized results from the ROSMAP. We found that the target genes of six of twenty-two TFs were significantly enriched in the AD excitatory neurons: RUNX1, YY1, HES1, SNAI2, PPARG and CEBPB (Figure 1C-D, Supplementary Figure 1B-E). Another key effect of TF activity is opening of their binding motifs, and Assay for Transposase-Accessible Chromatin through high-throughput Sequencing (ATAC-seq) represents a suitable tool to map regions of increased chromatin accessibility following TF binding ^43,44^. As we had previously generated ATAC-seq data from the same cohort of AD and control patients ^3^, we interrogated the data and searched for motifs with high binding affinity for these six TFs. We found that RUNX1 and YY1 motifs were significantly open in AD iNs (*q*-value < 0.0001) (Figure 1E).

Both RUNX1 and YY1 were substantially expressed and significantly upregulated in both AD iNs and excitatory neurons from human post-mortem AD brains (Figure 1F-G). Notably, their expression was specific to aged neurons (Figure 1F), as there was no significant differential expression in the isogenic induced pluripotent stem cell (iPSC)-derived iNs (*n* = 20) (Figure 1F), which reflect fetal neuron identities ^45–48^, or in aged fibroblasts from the same donors (Supplementary Figure 1F). Also, the expression of both RUNX1 and YY1 negatively correlated with the Mini-Mental State Examination (MMSE) obtained from the same patients, suggesting that high levels of these TFs in AD iNs correlated with poor cognitive functional scores (Supplementary Figure 1G-I).

These analyses captured two TFs, RUNX1 and YY1, that might act upstream of signaling pathways driving dysfunctional neuronal changes in AD.

### RUNX1 and YY1 overexpression in healthy aged iNs

To test if these putatively upstream-acting TFs contributed to increasing the susceptibility of aged neurons to develop cellular phenotypes such as neuronal identity loss and metabolic dysfunction associated with AD and a broader range of neurodegenerative disorders, we took advantage of our novel age-equivalent neuronal model (iNs). As women are more susceptible to developing AD, exhibit a more severe disease phenotype and account for the majority of AD cases, we selected a cohort of aged female donors for this study (Supplementary Table 1).

Using our established Ngn2/Ascl1-based iN conversion protocol ^3,17^, we generated two sets of iNs from aged but otherwise cognitively normal healthy donor fibroblasts and overexpressed either RUNX1 or YY1 independently, using a lentiviral system (Figure 2A-B, Supplementary Figure 2A). Following ∼19 days of neuronal conversion, iN cultures were transduced with either RUNX1 or YY1 and examined after eight days of transduction. Cultures for both TFs expressed NeuN and βIII-tubulin neuronal markers and green fluorescent protein marker (EGFP) (Figure 2C). Typical transduction efficiencies ranged from 25-97%, indicating that the TF overexpression was feasible and effective in the iNs (Figure 2D-E). To obtain near-pure transgenic neuron populations for downstream readouts, the cells were then isolated using FACS for PSA-NCAM+/EGFP+ double-positive cells (Figure 2D-E, Supplementary Figure 2B-G).

**Figure 2.**
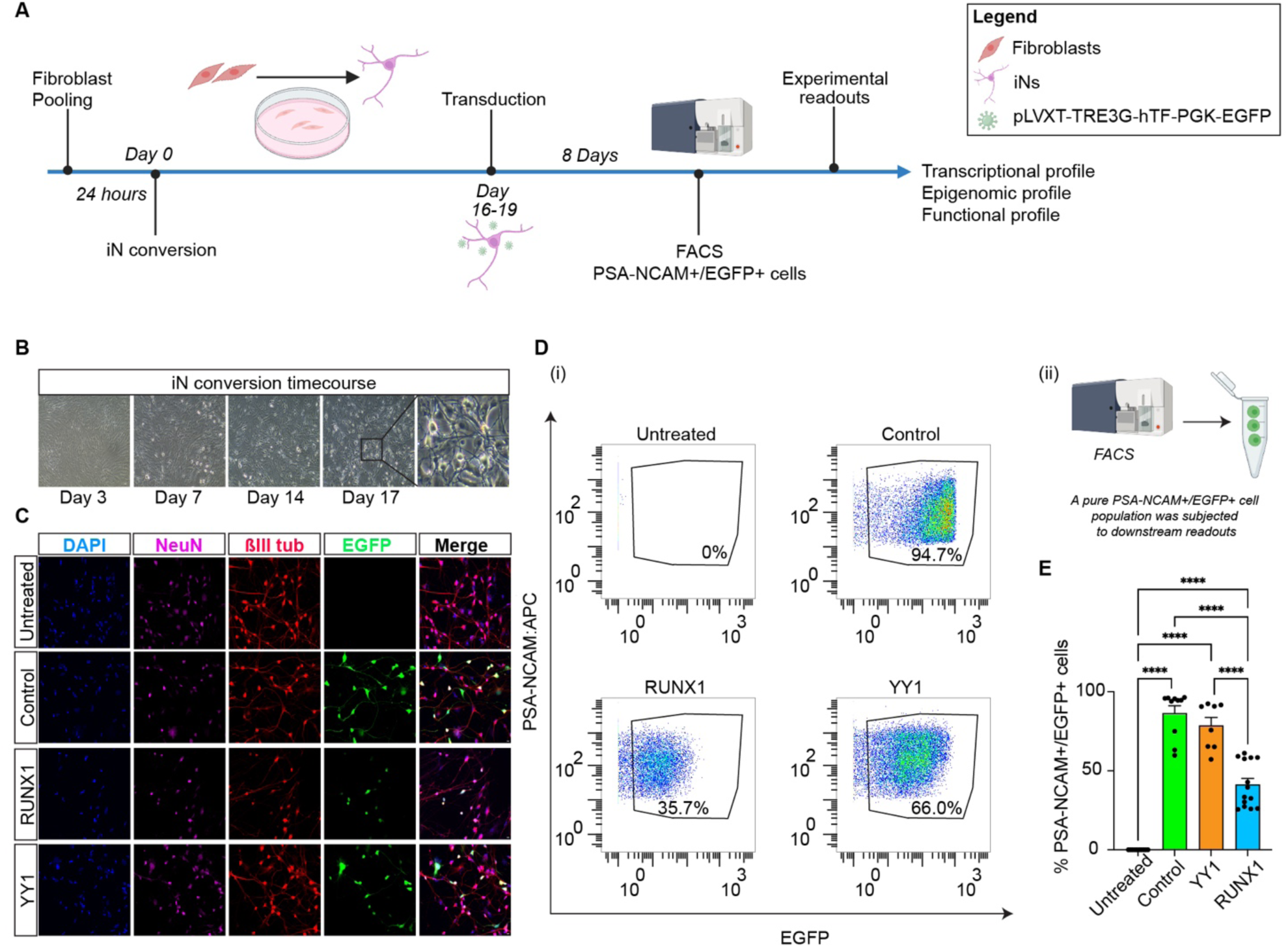
Gain-of-function study: RUNX1 and YY1 overexpression in aged iNs. (A) Schematics. Experimental design of the TF gain-of-function study in aged iNs. (B) iN conversion time course. Representative phase contrast images of fibroblast-to-neuron direct conversion on day 3 (high cell density) and at different stages of culture in NC media; scale bar, 10 μm (*n* = 3 and 3 independent iN culture replicates per donor). (C) Representative immunofluorescence images of iN cultures expressing neuronal markers, NeuN and βIII-tubulin, and green fluorescent protein, EGFP, 8 days post viral transduction; scale bar, 10 μm. (D) (i) Representative PSA-NCAM+/EGFP+ cells FACS gates. The iNs OE RUNX1 or YY1 were sorted by PSA-NCAM, neuronal marker, and by EGFP. Example plot and gating for FACS-based purification of untreated iNs (untreated), and iNs transduced with lentiviral particles containing plasmids carrying either the EGFP only (control), RUNX1 or YY1 sequence. (ii) Schematics. A pure population of PSA-NCAM+/EGFP+ double-positive cells was isolated using FACS for downstream readouts. (E) Percent of FACSorted PSA-NCAM+/EGFP+ cells over live single cells (*n* = 3 x Untreated, *n* = 3 x CTL, *n* = 3 x iNs OE YY1, *n* = 3 x iNs OE RUNX1, 1 to 5 independent iN culture replicates per condition). Statistical significance was assessed by performing one-way ANOVA, Tukey’s multiple comparisons test.

### RUNX1 overexpression promotes neuronal identity loss in aged iNs

To investigate the impact of the TFs on global neuronal gene expression, we performed RNA-seq. Principal Component Analysis (PCA) on all genes was conducted for dimensionality reduction. The analysis clearly distinguished between iNs OE RUNX1, YY1 and control iNs, indicating variation in gene expression profiles, with PC1 and 2 capturing the highest variance among the samples (Supplementary Figure 3A).

To evaluate the transcriptomic signature specifically induced by RUNX1, we calculated the differentially expressed (DE) genes between iNs OE RUNX1 and controls. Based on a stringent criterion (adj*p* value < 0.01, Log2FC > +1 or < -1), RUNX1 expression resulted in a substantial number of significant DE genes, with 981 significantly upregulated and 1,540 significantly downregulated genes (Figure 3A-B). RUNX1 transcript was highly significantly expressed in the iNs OE RUNX1, providing a further indication that the transduction of the gene in iNs was effective (Supplementary Figure 2B).

**Figure 3.**
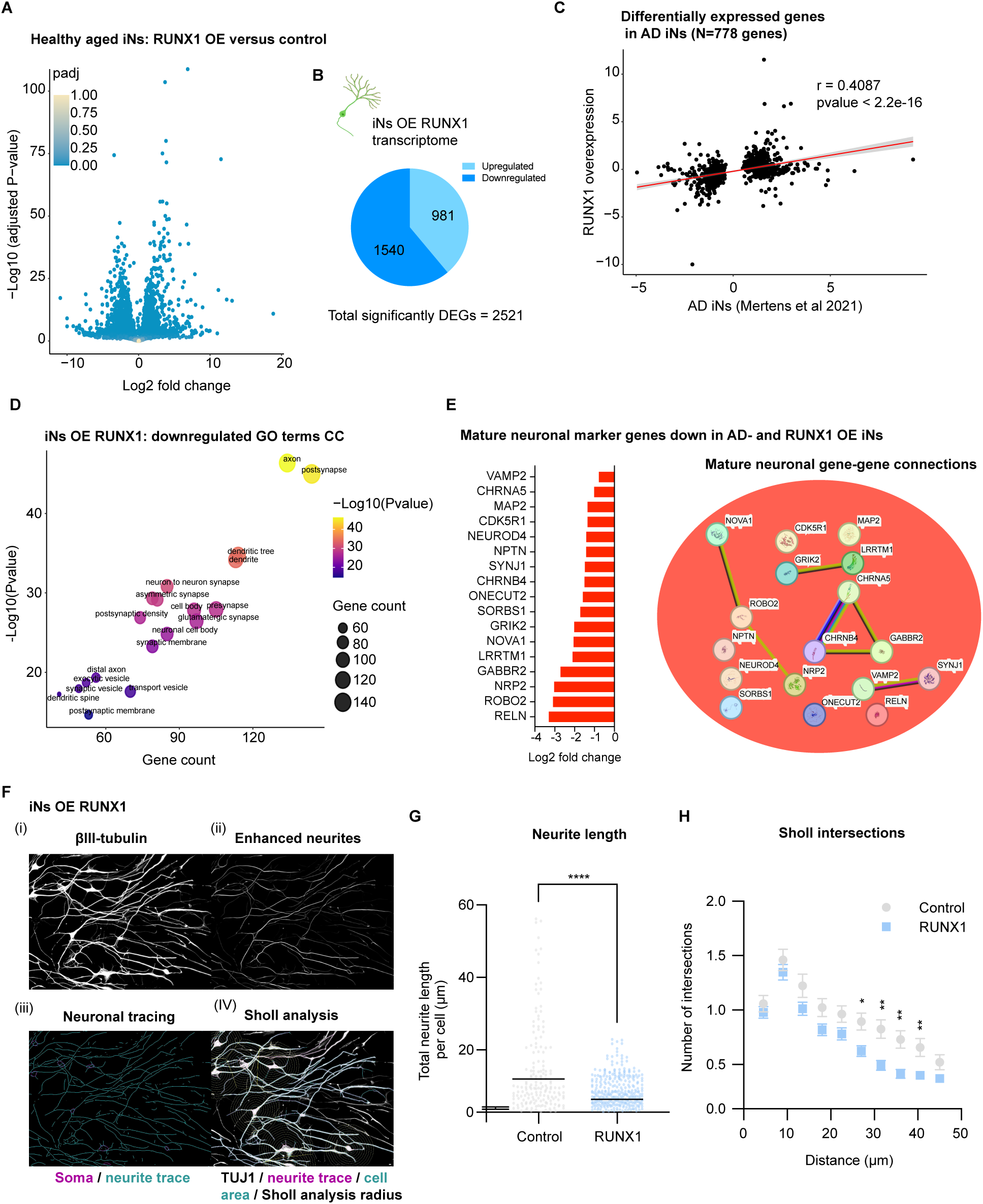
RUNX1 overexpression in aged iNs results in loss of cellular identity. (A) Volcano plot depicting DE genes from the iNs OE RUNX1 versus control (*n* = 3 versus 3 and 3 iN culture replicates per condition). (B) Total number of significantly up- and downregulated genes upon RUNX1 OE. (C) Pearson’s correlation between log2 fold-changes in AD iNs and in iN OE RUNX1 of genes that are differentially expressed in AD iNs (N=778) (^3^) on a gene-by-gene basis. (D) GO term analysis of significantly downregulated genes upon RUNX1 OE reveals specific loss of mature neuronal signature (dot plot illustrating top significantly depleted GO cellular components (CC) (Supplementary Table 2)). Point sizes correspond to the number of genes assigned to each category. (E) Bar graph showing log2 fold changes of downregulated mature neuronal marker genes in iNs OE RUNX1 versus control (left panel). STRING network showing gene-gene connections across the represented downregulated neuronal markers (right panel). (F) Neuronal morphometric reconstruction of iNs OE RUNX1. (i) Representative immunofluorescence image of iNs OE RUNX1 stained for neuronal marker βIII-tubulin, (ii-iV) neuronal reconstructions and Sholl analysis using Cell Profiler. Scale bar 10 μm. Experiments were performed in all 3 patient lines and 2 iN culture replicates per condition. Results are shown in G and H. (G) The iNs OE RUNX1 showed reduced growth properties compared to control (each dot represents a cell; statistical relevance was assessed using parametric unpaired *t*-test, \*\*\*\**p*<0.0001; values show means ± S.E.M.). (F) Sholl analysis of branching patterns between iNs OE RUNX1 and control. Sholl neurite intersections were reduced in the iNs OE RUNX1 as compared with controls (statistical significance was assessed using two-way ANOVA with Sidak correction, *adjusted *p*<0.05, **adjusted *p*<0.01; values represent mean ± S.E.M.).

We first examined how these gene expression changes correlated with the changes observed in AD iNs. Specifically, we compared the expression of the significantly DE genes from AD iNs with the expression of the same genes in the RUNX1 dataset; results showed a significant positive correlation between the two datasets (Figure 3C). To examine the biological relevance of all genes significantly up- and downregulated by RUNX1 OE, we performed gene ontology (GO) analysis. While the effect on the upregulated genes was less clear, yielding a heterogeneous set of terms with genes associated with different pathways (Supplementary Figure 3D), the response of the downregulated genes was evident. Strikingly, the expression of just RUNX1 in healthy aged iNs was characterized by depletion of synaptic genes and other genes promoting mature structural and functional neuronal identity, including axon, post-synapse and dendritic tree GO terms, among the cellular components, as well as biological processes including neuron projection development, modulation of chemical synaptic transmission and cell morphogenesis (Figure 3D, Supplementary Figure 3E and Supplementary Table 2). These changes accounted for genes like MAP2 and MAP1B (Supplementary Figure 3C), whose loss of function is a feature of a vulnerable neuronal state and is tightly linked to the neuronal dysfunction seen in early AD neurodegeneration ^49–52^. We also tested a curated list of previously defined mature neuronal markers that are also highly significantly downregulated in AD iNs ^3,53^. Results showed that these genes were markedly downregulated following RUNX1 overexpression and highlighted loss of genes with a strong synaptic and structural integrity relevance, including RELN, VAMP2 and NEUROD4 (Figure 3E), which are strongly implicated in entorhinal cortex degeneration ^54–59^.

We and others have previously shown that loss of a mature neuronal transcriptional profile underlies neuronal identity instability in AD and reduced neuronal arborization complexity is a strong indicator of this instability. To query if these changes in expression impacted intrinsic maturational properties, we examined neuronal arborization complexity. Notably, and consistent with changes observed in AD iNs ^3^, systematic reconstruction of individual morphological neuronal tracing revealed a significant reduction in neuronal arborization complexity in iN OE RUNX1 and control cultures (Figure 3F-G, Supplementary Figure 3F). Additional measures indicated that the cells featured branching patterns that were significantly less complex than those of control iNs (Figure 3H).

Altogether the data show that high levels of RUNX1 result in the downregulation of neuronal identity genes that is sufficient to produce AD-like structural deficiencies in otherwise healthy neurons.

### YY1 overexpression modulates changes in energy metabolism in aged neurons

Next, we turned our attention to our other candidate TF. A parallel analysis was performed on RNA-seq data generated from the iNs OE YY1. Induction of YY1 in iNs resulted in large changes in gene expression, consisting of 691 significantly upregulated and 483 downregulated genes (adj*p* value < 0.01, Log2FC > +1 or < -1) (Figure 4A-B), indicating a strong regulatory effect of YY1 on the transcriptome of adult neurons. The YY1 transcript was highly significantly expressed in the iNs OE YY1, providing a further indication that the transduction of this gene in iNs was effective (Supplementary Figure 4A). Correlation analysis between the expression of AD iN significantly expressed genes and their changes in iN OE YY1 showed a mild but highly significant positive correlation between the AD iN signature and the YY1-induced transcriptional profile (Figure 4C).

**Figure 4.**
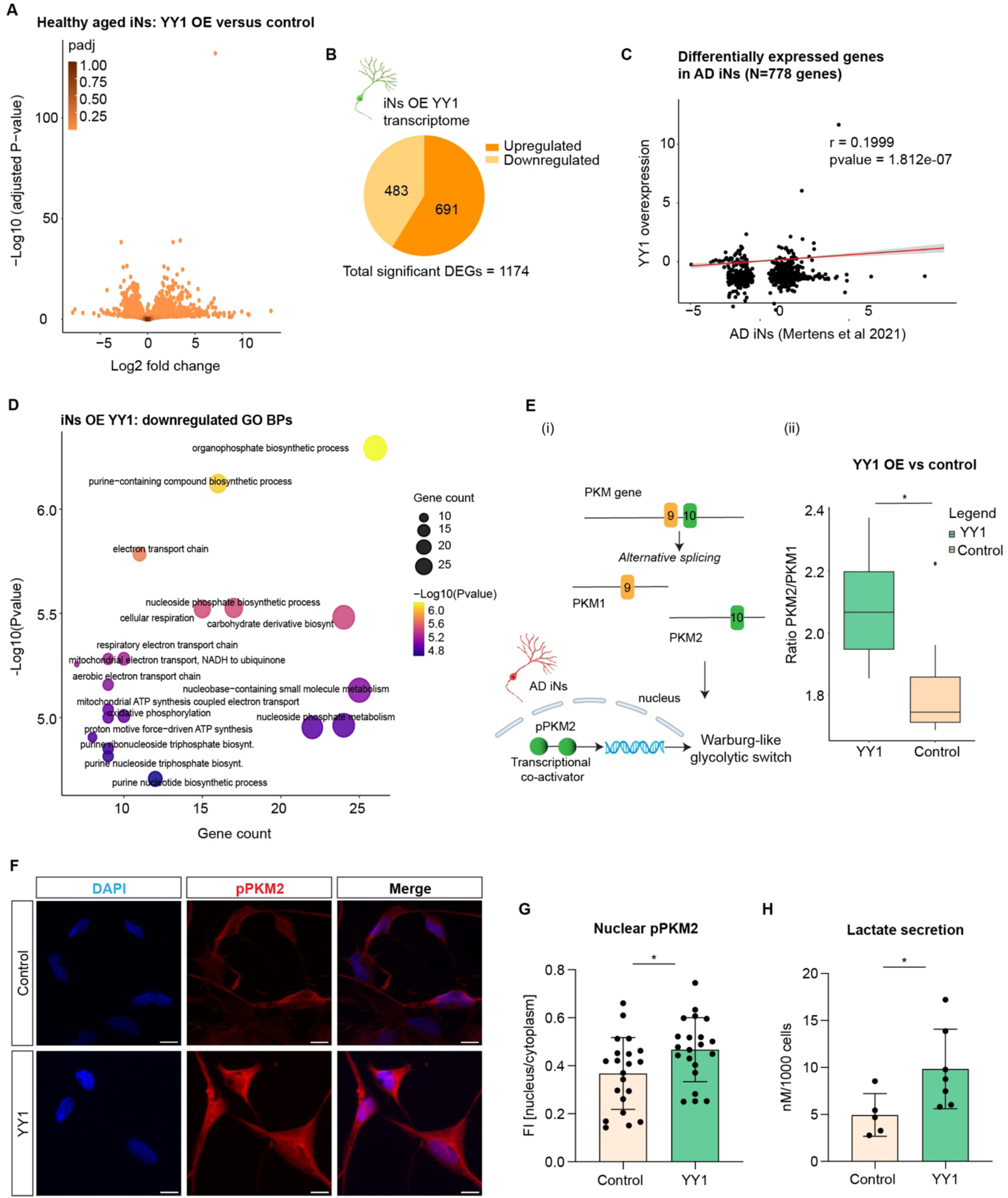
YY1 overexpression in aged iNs drives transcriptional reprogramming of neuronal energy metabolism. (A) Volcano plot showing DE genes from the iNs OE YY1 versus control. B) Total number of significantly up- and downregulated genes upon YY1 OE. (C) Pearson’s correlation between log2 fold-changes in AD iNs and in iN OE YY1 of genes that are differentially expressed in AD iNs (N=778) (^3^) on a gene-by-gene basis. (D) GO term analysis of significantly downregulated genes upon YY1 OE reveals loss of mature energy metabolism gene programs (dot plot showing top significantly depleted GO biological processes (BPs) (Supplementary Table 3)). Point sizes correspond to the number of genes assigned to each category. (E) Schematics of the PKM gene highlighting exon 9 and 10 and of the PKM alternative splicing generating PKM1 and PKM2 isoforms, depending on exon retained; nuclear pPKM2 translocation in AD iNs drives Warburg-like metabolic switch (i) (^7^); PKM2/PKM1 expression ratio increases in the iNs OE YY1 compared to control (ii). (F-G) Immunostaining (F) and quantification (G) of pPKM2(Ser37) (*n* = 4 control iNs, *n* = 2 iNs OE YY1, and 1 to 8 independent iN culture replicates per condition; each dot represents one image; significance, unpaired *t*-test). Scale bars, 10 μm. (H) Colorimetric assay was performed to measure secreted lactate in the supernatant of the iNs OE YY1 and control (*n* = 3 control iNs, *n* = 2 iNs OE YY1 and 1 to 4 independent iN culture replicates per condition; each dot represents one independent iN culture; significance, unpaired *t*-test).

Functional enrichment analysis was then performed on YY1-induced DE genes. As we had seen with RUNX1, while upregulated genes appeared functionally diffuse and did not suggest a clear underlying effect (Supplementary Figure 4B and Supplementary Table 3), the effect on the downregulated genes was more specific. GO analysis showed clear loss in mature energy metabolism-related gene sets (Figure 4D, Supplementary Figure 4C-E and Table 3) including nucleotide metabolism, cellular respiration and oxidative phosphorylation, which is the major energy source for adult neurons ^7,23^.

We have previously shown that AD iNs undergo a Warburg-like metabolic shift towards aberrant glycolysis, which occurs through the isoform switch from glycolytic enzyme pyruvate kinase M (PKM1) to the cancer-associated isoform PKM2. Specifically, in AD iNs, we observed an increase in the PKM2/PKM1 ratio and an increased nuclear translocation of phosphoPKM2 (Ser37) (pPKM2), which ultimately mediates the observed metabolic reprogramming (Figure 4E (i)) ^23^. To examine if the transcriptional changes driven by YY1 resulted in any of these functional alterations, we first measured the PKM2/PKM1 ratio and found that this ratio was significantly higher in the iNs OE YY1 (Figure 4E). Then we examined nuclear translocation of pPKM2 through immunohistochemistry; results showed an increased nuclear translocation of pPKM2 following YY1 overexpression (Figure 4F-G and Supplementary Figure 4F), consistent with increased lactate secretion (Figure 4H and Supplementary Figure 4G), which is indicative of increased glycolysis.

Taken together, these data indicate that YY1 overexpression in aged iNs reshapes the transcriptional energy profile of the cells causing specific functional changes causative of metabolic dysfunction in AD neurons.

### RUNX1 and YY1 overexpression enhances accessibility at *cis*-regulatory sites associated with AD-related signaling pathways

To map differential *cis*-regulatory chromatin accessibility driven by RUNX1 and YY1 OE in iNs, we next subjected PSA-NCAM+/EGFP+ cells to ATAC-seq ^43,44^. This analysis enabled accurate mapping of the genomic landscape of the iNs and revealed a profound chromatin remodeling upon TF overexpression. Specifically, we identified 30,912 uniquely significant (*p* < 0.0001) opened and 11,687 closed ATAC-seq peaks upon RUNX1 overexpression (Figure 5A-B) and 12,429 significantly open and 3,228 significantly closed peaks upon YY1 overexpression (Figure 5C-D).

**Figure 5.**
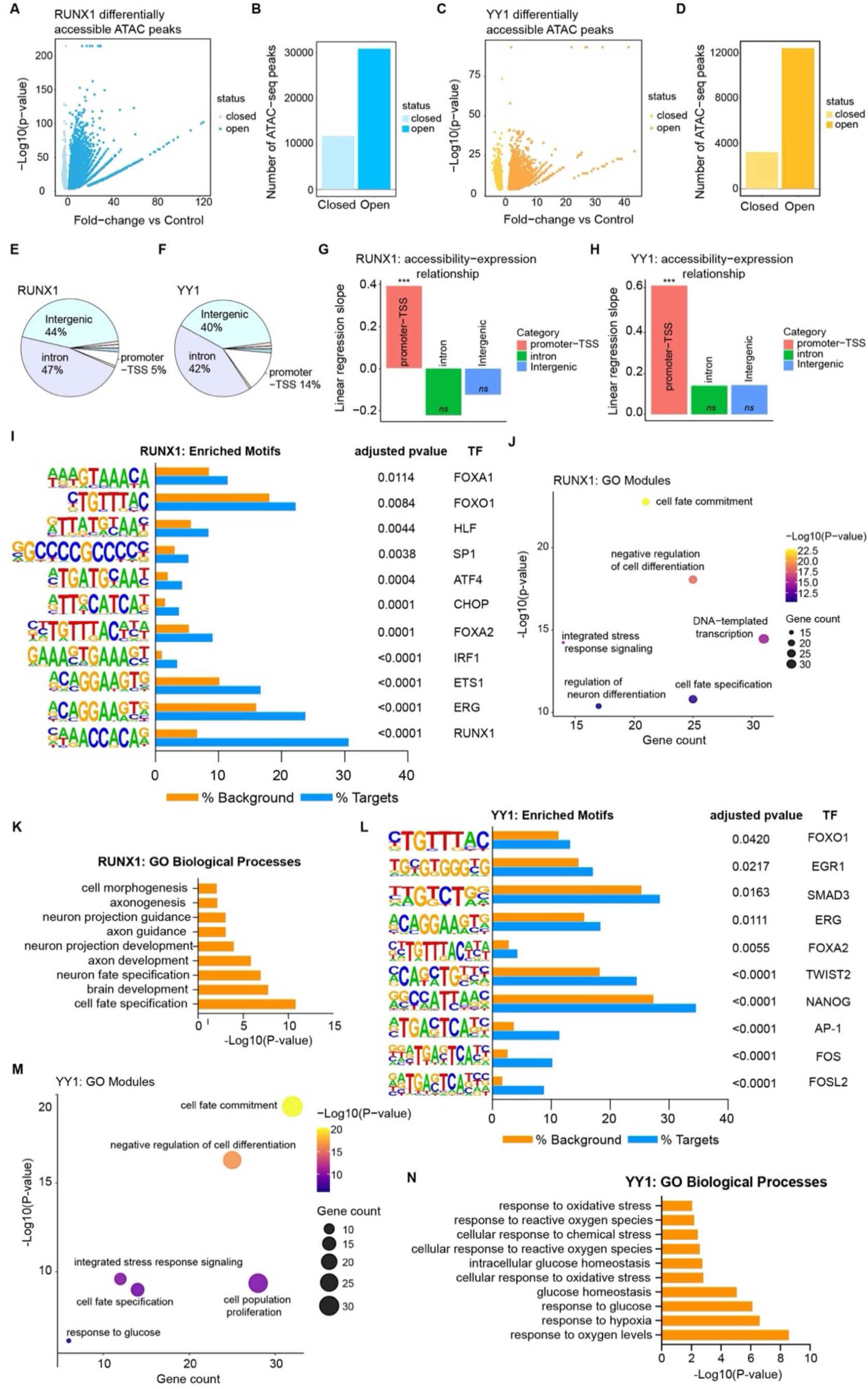
ATAC-seq reveals a dynamic opening of *cis*-regulatory landscape associated to AD neurodegeneration in iNs OE RUNX1 and YY1. (A-D) Volcano plot and quantification of differentially accessible ATAC-seq peaks in the iNs OE RUNX1 OE versus control, respectively (A-B) and in the iNs OE YY1 (C-D); each dot on the volcano plots represents an ATAC-seq differential peak (*n* = 4 x iN controls, *n* = 4 x iNs OE RUNX1 and *n* = 4 iNs OE YY1; *p* < 0.0001 and fold change | >1|). (E-F) Pie charts quantifying the distribution of ATAC-seq peaks per genomic annotation, in the RUNX1 OE iNs (E) and YY OE iNs (F), respectively. (G-H) Bar plots displaying linear regression slopes between ATAC-seq peak accessibility and RNA-seq expression for promoter-TSS, intronic, and intergenic differential peaks in the RUNX1 OE iNs (G) and YY OE iNs (H), respectively. Asterisks indicate significant regression (\**p* < 0.05, \*\**p* < 0.01 and ***p < 0.001. *P*-values for RUNX1 was 2.01e-14; *p*-value for YY1 < 2e-16). (I) HOMER motif enrichment analysis of differentially open ATAC-seq peaks from the iNs OE RUNX1 (Supplementary Table 4). Bar graph for exemplified motifs showing percentages of target and background sequences with motif, *q* values for motif enrichment and corresponding high-affinity DNA-binding TF. (J-K) GO analysis of the TFs with high binding affinity to all significantly enriched motifs within the promoter-TSS in the iNs OE RUNX1. Bubble plots for BPs displaying selected BP modules (i.e. cluster terms grouping GO terms based on their semantic similarity using Metascape); point sizes correspond to the number of genes within to each cluster and they are colored by significance (J and Supplementary Table 5); bar graph shows selected significantly enriched GO BPs (K and Supplementary Table 5). (L) HOMER motif enrichment analysis of differentially open ATAC-seq peaks from the iNs OE YY1 (Supplementary Table 6). Similarly to (I), bar graph illustrates exemplified motifs with percentages of target and background sequences with motif, *q* values for motif enrichment and corresponding high-affinity DNA-binding TF. (M-N) GO analysis of the TFs with high binding affinity to all significantly enriched motifs within the promoter-TSS in the iNs OE YY1. Bubble plots for BPs displaying selected BP modules; point sizes correspond to the number of genes within to each cluster, colored by significance (M and Supplementary Table 7); bar graph shows selected significantly enriched GO BPs (N and Supplementary Table 7).

In iN OE RUNX1, the majority of the significant ATAC-seq peaks (47%) were located within intronic regions, 44% in the intergenic regions and 5% within the promoter-Transcription Start Site (TSS) (Figure 5E). Pairing these data with gene expression from RNA-seq data showed that changes in chromatin accessibility within the promoter-TSS significantly correlated with changes in gene expression (Figure 5G). Similarly, in iN OE YY1, 42% of the significant ATAC-seq peaks, which accounted for the majority, were located within the intronic regions, 40% were located in the intergenic regions and 14% within the promoter-TSS (Figure 5F). Changes in chromatin accessibility at promoter-TSS correlated with gene expression changes (Figure 5H). Overall, the chromatin accessibility profile induced by RUNX1 and YY1 revealed a global shift towards a more accessible chromatin state, which is structurally more permissive to regulatory factor binding.

To gain insights into the gene regulatory landscape most affected by the chromatin changes, we performed short-sequence motif enrichment analysis of promoter-TSS open peaks using HOMER and matched the motifs against known TF binding databases. Motif discovery analysis of the significantly open peaks within the promoter-TSS in iNs OE RUNX1 revealed significant enrichment for 119 short-sequence binding motifs (Supplementary Table 4 and Supplementary Figure 5A). The most enriched motif in opening regions binds RUNX1 (Figure 5I), indicative of an active functional role for RUNX1 when expressed in aged iNs. Motifs with high binding affinity for regulators of cell fate commitment became more accessible, including FOXO1, HLF and ERG1 (Figure 5I) ^60–66^, which are TFs associated with cellular de-differentiation and identity instability in aging and AD ^6^, as well as immediate early genes, such as AP-1 and FOS (Supplementary Figure 5A) ^67–70^. Of note, some of the top motifs corresponded to high-affinity sites for key pioneer factors, such as the FOXA1/2 TF family, that can penetrate and open highly compacted DNA structures, rendering them more accessible to TF binding (Figure 5I). Interestingly, other TF motifs previously observed in AD neurons ^3,8^ were also enriched in iNs OE RUNX1. For instance, data revealed a significant overrepresentation of binding sites for ATF4 and CHOP, two key Immediate Stress Response (IRS) TFs (Figure 5I) ^71–74^. More accessibility for SP1, a key regulator involved in the cellular response to nitrogen compounds, a process linked to genomic instability, was also observed (Figure 5I) ^75–77^. In addition, the ETS1 motif, a potential key upstream regulator of neuronal senescence gene expression, was significantly more open in iNs OE RUNX1 (Figure 5I) ^78–80^. TFs with elevated activity in significantly enriched *cis*-regulatory elements were enriched for regulatory networks that control cell fate specification, chromatin remodeling, cancer and senescence as shown by GO and Kyoto Encyclopedia of Genes and Genomes (KEGG) pathway analysis (Figure 5J-K, Supplementary Figure 5B-C and Supplementary Table 5).

On the other hand, HOMER motif discovery analysis of differentially promoter-TSS open peaks in iN OE YY1 identified 74 significantly over-represented motifs (Supplementary Table 6) corresponding to high binding affinity sites for several TFs, including some previously probed in AD iNs, for instance AP1 and FOS, involved in genomic instability ^64,81–90^. Strikingly, open motifs were also found in immediate early genes including EGR1, whose dysfunction can be associated with metabolic drift and neuropathology in AD neurons ^91–93^ (Figure 5L and Supplementary Figure 5D). Other TFs with high affinity for differentially open motifs included ATF3 and BACH1, which are also involved in the IRS (Supplementary Figure 5D) ^94–98^. Functional enrichment analysis of all the TFs with elevated activity in significantly gained promoter-TSS peaks showed their involvement in networks controlling cell differentiation, response to glucose and response to hypoxia, among others (Figure 5M-N, Supplementary Figure 5E-F and Supplementary Table 7).

These results highlight that RUNX1 and YY1 overexpression induces an extensive reshaping of the chromatin landscape, characterized by a large increase in chromatin accessibility, and increases accessibility in *cis*-regulatory elements for TF networks driving signaling pathways associated with increased neuronal vulnerability in AD, reprogramming the chromatin landscape of aged neurons toward an AD-like, degeneration-prone state and predisposing neurons to transcriptional collapse.

### Downregulating RUNX1 and YY1 expression ameliorates neuronal features in AD

To evaluate whether suppression of RUNX1 and YY1 activity would be sufficient to restore some features of the mature neuronal phenotype in AD iNs and identify them as therapeutical targets, we next performed a loss-of-function experiment (Figure 6A). We generated iNs directly from AD donor fibroblasts and expressed short hairpin (sh)RNAs to knockdown (KD) RUNX1 or YY1, respectively. We designed constructs for three distinct shRNAs targeting RUNX1 or YY1 (Supplementary Figure 6A) and selected the shRNA that achieved the most efficient target KD, exemplified by RUNX1 shRNA1 and YY1 shRNA1 with ∼63% and 72% RUNX1 and YY1 KD efficiency, respectively (Supplementary Figure 6B-C). Following iN conversion and ∼8 days of shRNA transduction, AD iN cultures expressing either scramble (SCRM), RUNX1 or YY1 were subsequently isolated using FACS (Supplementary Figure 6E-J). FACS isolation resulted in cultures of over 80% of mCherry-positive neuronal cultures across all the conditions, confirming efficient delivery of the shRNA (Supplementary Figure 6J). These data provided evidence that the shRNA strategy to modulate gene expression is feasible and effective in human iNs.

**Figure 6.**
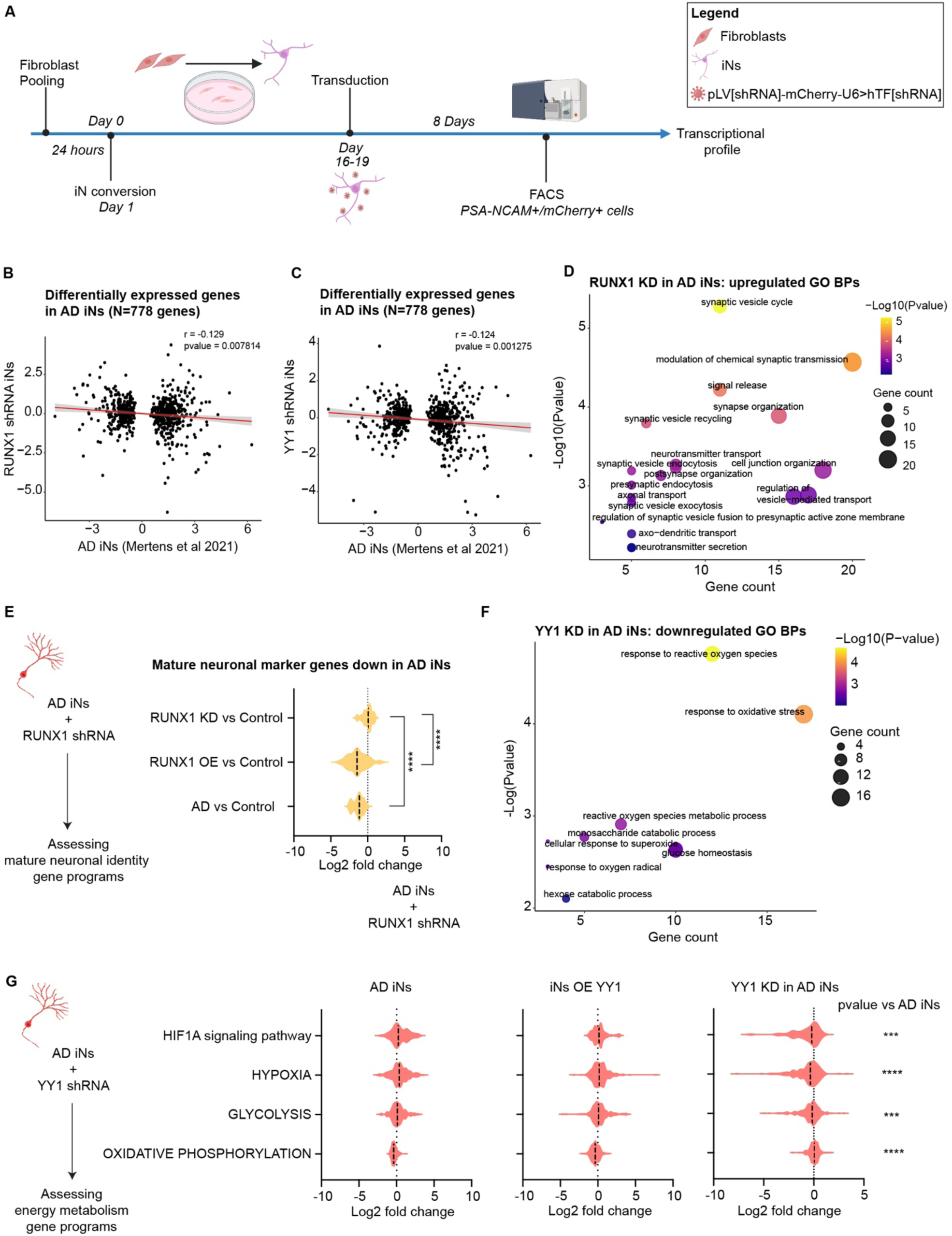
Downregulation of RUNX1 and YY1 in AD iNs reinstates gene regulatory programs associated with a healthy mature neuronal phenotype. (A) Schematics. Experimental design of the TF loss-of-function study in AD iNs. (B-C) Pearson’s correlation between AD iN and the AD iN transcriptome upon RUNX1 KD (B) and YY1 KD (C) of log2 fold-changes of genes that are differentially expressed in AD iNs (N=778) (^3^) on a gene-by-gene basis. (D) GO analysis of upregulated genes in AD iNs upon RUNX1 KD reveals upregulation of genes promoting mature neuronal identity. Bubble plots for selected biological (Supplementary Table 8); point sizes correspond to the number of genes within to each cluster, colored by significance. (E) Violin plot showing expression changes of mature neuronal marker genes in the AD iNs, iNs OE RUNX1 and AD iNs upon RUNX1 KD. The expression pattern of these genes in aged iNs mirrors that observed in AD iNs following RUNX1 OE, and it is reversed when RUNX1 is downregulated in AD iNs. Statistical significance was assessed using Wilcoxon test (**** *p*<0.0001). (F) GO analysis of downregulated genes in AD iNs following YY1 KD identifies downregulation of pathways related to glucose metabolism and oxidative stress response, indicating a shift in transcriptional programs driving AD-associated neuronal metabolic phenotype. Bubble plots for selected biological (Supplementary Table 9); point sizes correspond to the number of genes within to each cluster, colored by significance. (G) Violin plots showing expression changes in iNs OE YY1 versus control, and in AD iNs following YY1 KD, of genes involved in signaling pathways dysregulated in AD. Statistical significance was assessed using Wilcoxon test (*** *p*<0.001; **** *p*<0.0001).

As the cellular transcriptome encodes the instructions specifying cell state and fate, we chose RNA-seq as our primary readout. Therefore, RNA was extracted from FACS-purified PSA-NCAM+/mCherry+ AD iNs and subjected to RNA-seq. Using PCA dimensionality reduction including all genes, RUNX1 and YY1 KD induced two largely distinct transcriptional profiles clustering separately from the AD iNs, with PC2 and 3 capturing the highest variance among the samples (Supplementary Figure 6D). Correlation between the AD iNs and either RUNX1 KD-induced or YY1 KD-induced gene expression profile showed a significant negative correlation between the two datasets (Figure 6B-C), indicating that RUNX1 and YY1 downregulation reverses aspects of the AD-associated transcriptional profile.

GO analysis of the DE genes induced by RUNX1 KD in AD iNs revealed highly significant enrichment for genes controlling specific aspects of mature neuronal identity such as signal release from synapse, axonal transport, synapse organization and cellular homeostasis, as well as neuronal components such as axon, neuron projection cytoplasm and dendrite (Figure 6D and Supplementary Table 8). We also tested the expression of a curated list of mature neuronal markers significantly downregulated in AD iNs ^3^ and iN OE RUNX1, and we found that their expression was ameliorated upon RUNX1 KD (Figure 6E).

On the other hand, GO term analysis of the DE genes induced by YY1 KD in AD iNs showed enrichment of genes involved in energy metabolism, revealing a significant downregulation of pathways, including glucose metabolism and oxidative stress (Figure 6F and Supplementary Table 9). We also tested the expression of genes involved in major pathways disrupted in AD neurons and iNs OE YY1, such as OXPHOS and glycolysis, and data showed that the expression of genes involved in these pathways significantly changed upon YY1 KD compared to AD iNs and iN OE YY1 (Figure 6G).

Altogether, these data indicate that RUNX1 and YY1 downregulation in AD iNs reinstates gene regulatory programs associated with a healthy mature neuronal phenotype, ameliorating complementary aspects of the AD iN phenotype.

## Discussion

This study identifies RUNX1 and YY1 as age-associated transcriptional master regulators that orchestrate loss of neuronal fate and metabolic dysfunction in AD neurons, and it shows that interfering with these factors restores regulatory programs associated with a healthy neuronal phenotype. Loss of cell fate and metabolic dysfunction are in part responsible for neuronal impairment in AD ^2,3,23,99,100^. Previous post-mortem brain and *in vitro* studies have shown that AD neurons activate early developmental and cancer-associated genes involved in cell fate determination and energy metabolism putatively as an IRS ^1–3,5–8,15,23^. However, as in the case of other diseases, if sustained, expression of these genes can promote irreversible state and fate changes that lead to neurodegeneration ^6,7,23,101–104^. In this context, we found that AD neurons overexpress RUNX1 and YY1, two early developmental and cancer-related genes, and our current work suggests that they act as major age-associated modulators of the AD neuronal phenotype by contributing to increasing the susceptibility of the cells toward two major AD features, cell identity loss and metabolic dysfunction, respectively.

Our data position RUNX1 as an aging context-dependent driver of neuronal de-differentiation in AD. In epigenetically old human iNs, RUNX1 overexpression represses mature neuronal gene programs, leads to arborization defects, reducing morphological complexity, and opens genomic motifs with high binding affinity for early development, cell fate commitment TFs and stress- and senescence-associated regulatory elements. Furthermore, RUNX1 KD in AD iNs restores neuronal identity markers and partially normalizes neuronal transcriptional profile. These findings extend prior work implicating RUNX1 in hematopoietic and neuronal lineage specification ^26,29,31–38,105^. In the hippocampal precursor cells, RUNX1 expression is upregulated by TGF-β1 and Notch1 and decreases upon differentiation ^106,107^. In the adult brain, RUNX1 is expressed at low levels; however, its expression increases in response to traumatic injury and repair ^28,39^ , and our data add the notion that, in the aging brain context, aberrant RUNX1 re-activation can shift neurons back toward an immature, vulnerability-prone state. Also, RUNX1 is considered a cancer-related gene, in part because mutations in RUNX1 can lead to leukemia ^108–110^. Interestingly, RUNX1 is located on the long arm of chromosome 21 and has been associated with Down Syndrome (DS), where, along with other genes, it has a genomic dosage-dependent effect and can alter neuronal identity ^31,111–114^. Of note, it has been reported that individuals with DS are exposed to an increased risk of dementia ^3,115^, pointing towards an association between genes on chromosome 21 and increased susceptibility to dementia. In addition, a recent human postmortem brain study from AD individuals has identified RUNX1 as one of the major TFs enriched in the enhancers of AD-risk loci in microglia suggesting an active regulatory role for this TF in the context of AD ^8^. However, little is known about how this gene is involved in neurodegeneration at the cellular and molecular levels. Our work reveals that RUNX1 is expressed in AD neurons and shows that, when expressed in aged neurons, increases their vulnerability toward those cellular phenotypes, particularly loss of neuronal fate, that lead to AD-associated neuronal immaturity. As in the case of other cellular contexts, re-activation of RUNX1 in AD neurons might be a stress-associated response. Our data show that RUNX1-induced changes are part of the homeostatic response of the neuronal populations to the neurodegeneration described in AD brains ^2^ that ultimately drive neuronal vulnerability and neurodegeneration.

In parallel, our work identified YY1 as a second developmental and cancer-associated TF that is aberrantly engaged in AD neurons, reshaping gene regulatory programs that control energy metabolism. YY1 is a highly conservative zinc-finger TF that is ubiquitously expressed and whose function varies in a context-dependent manner ^116–120^. During mammalian embryogenesis, it modulates the expression of genes controlling neuronal lineage specification and migration ^117,119^ and promotes cerebellar cortical neural progenitor cell differentiation by suppressing the sex-determining region Y-box 2 expression ^121^. In human neural progenitor cells, it facilitates the progenitor-to-mature neuron transition by contributing to the regulation of LINE1 expression ^25^. YY1 also plays an important role in cellular bioenergetics during development ^40^. During early stages of mouse corticogenesis, YY1 exhibits a key role in the transcriptional regulation of mitochondrial bioenergetics and protein synthesis to guarantee neural progenitor cell proliferation and survival ^40^. Intriguingly, an *in silico* analysis from a previous post-mortem AD human brain studies showed that increased levels of YY1 mRNA have been found in human postmortem brain tissue and in laser-captured neurons from subjects with AD compared to controls, pointing towards YY1 as having an active regulatory role in AD ^42^. Notably, under pathological conditions, YY1 is re-activated and plays a major role in reprogramming cellular metabolism. In satellite muscle stem cells, during regeneration upon injury, YY1 regulates the metabolic switch towards glycolysis by directly downregulating mitochondrial pathway-related genes and upregulating glycolytic genes by stabilizing HIF1A ^24^. Of note, AD neurons undergo a similar Warburg-like metabolic rewiring, which involves HIF1A ^23^, and YY1 might contribute to stabilizing HIF1A in AD neurons. In this regard, a previous study showed that YY1 promotes the Warburg effect in cancer cells ^41^. Consistent with these findings, our data highlighted a conserved role for YY1 in coordinating stress-induced bioenergetic adaptation and revealed that, in AD iNs, YY1 reshapes energy metabolism regulatory programs by regulating the expression of genes perturbed in major signaling pathways such as OXPHOS, glycolysis and hypoxia that underlie neuronal metabolic dysfunctions in AD.

Here, we have identified and elucidated the implication of two TFs, RUNX1 and YY1, in the biology of the neuronal pathogenesis of AD. We have provided a mechanistic foundation to study their role as putative therapeutical targets for treatment of AD and age-associated neurodegeneration. Because RUNX1 and YY1 lie upstream of broad regulatory chromatin and transcriptional programs associated with AD dysfunctional changes, they may represent more efficient intervention targets than downstream factors. While we tested the specific contribution of these genes to the AD neuronal phenotype, individually, which is essential to probe their mechanistic role in the disease pathogenesis, future studies are needed to assess the interplay between the two and the impact on the onset and severity of trajectories driven by these TFs.

In this study, we have developed a novel approach to perturb age-associated genes in iNs and assess their contribution to cellular phenotypes implicated in AD. Given the growing interest in probing the role of genes identified from multi-omics atlases, in neurodegeneration, our methodological framework provides a valuable platform to be explored for future mechanistic studies aimed at identifying genes that contribute to AD and age-associated neurodegenerative disorder risk.

## Methods

### Key resource table

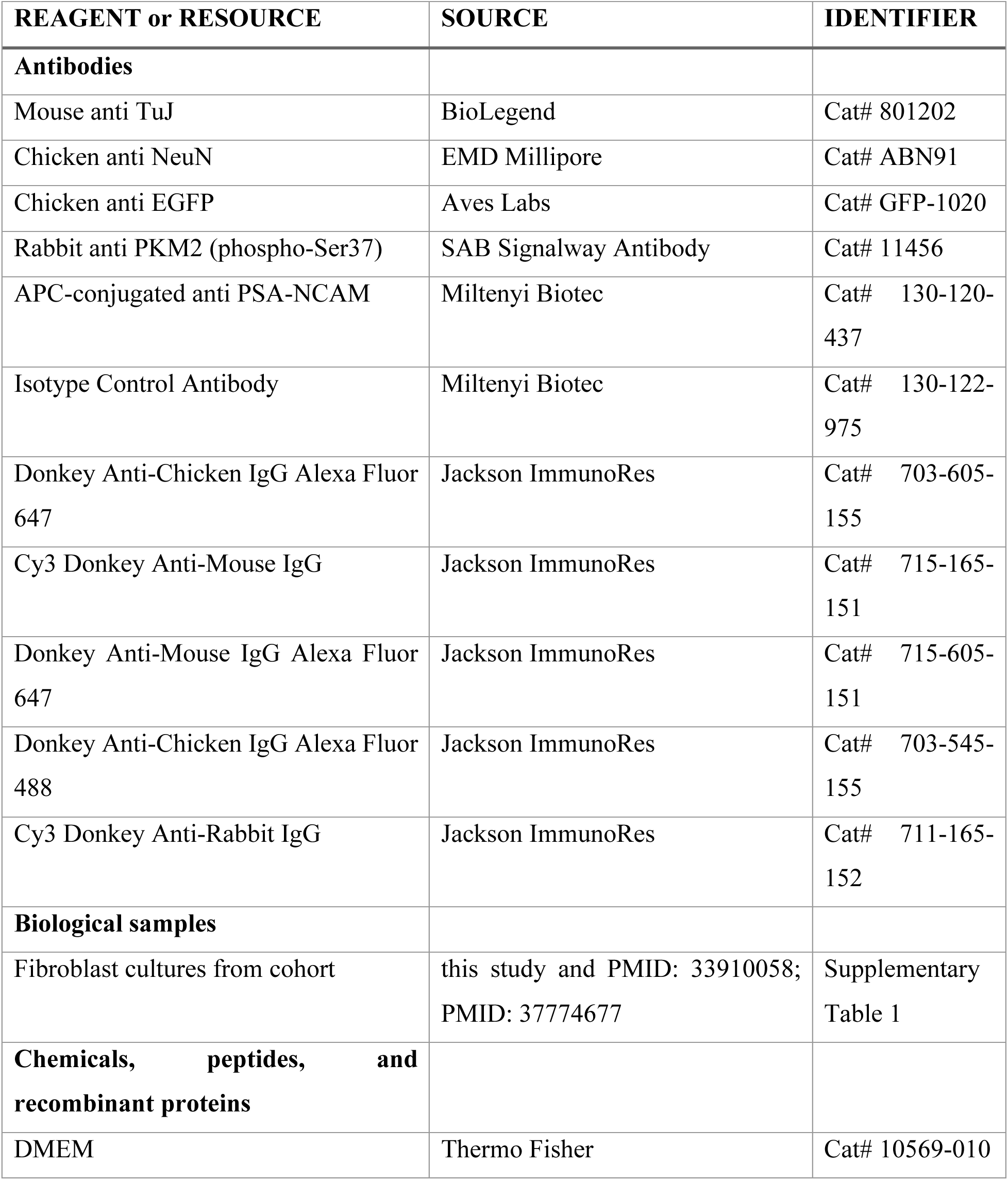
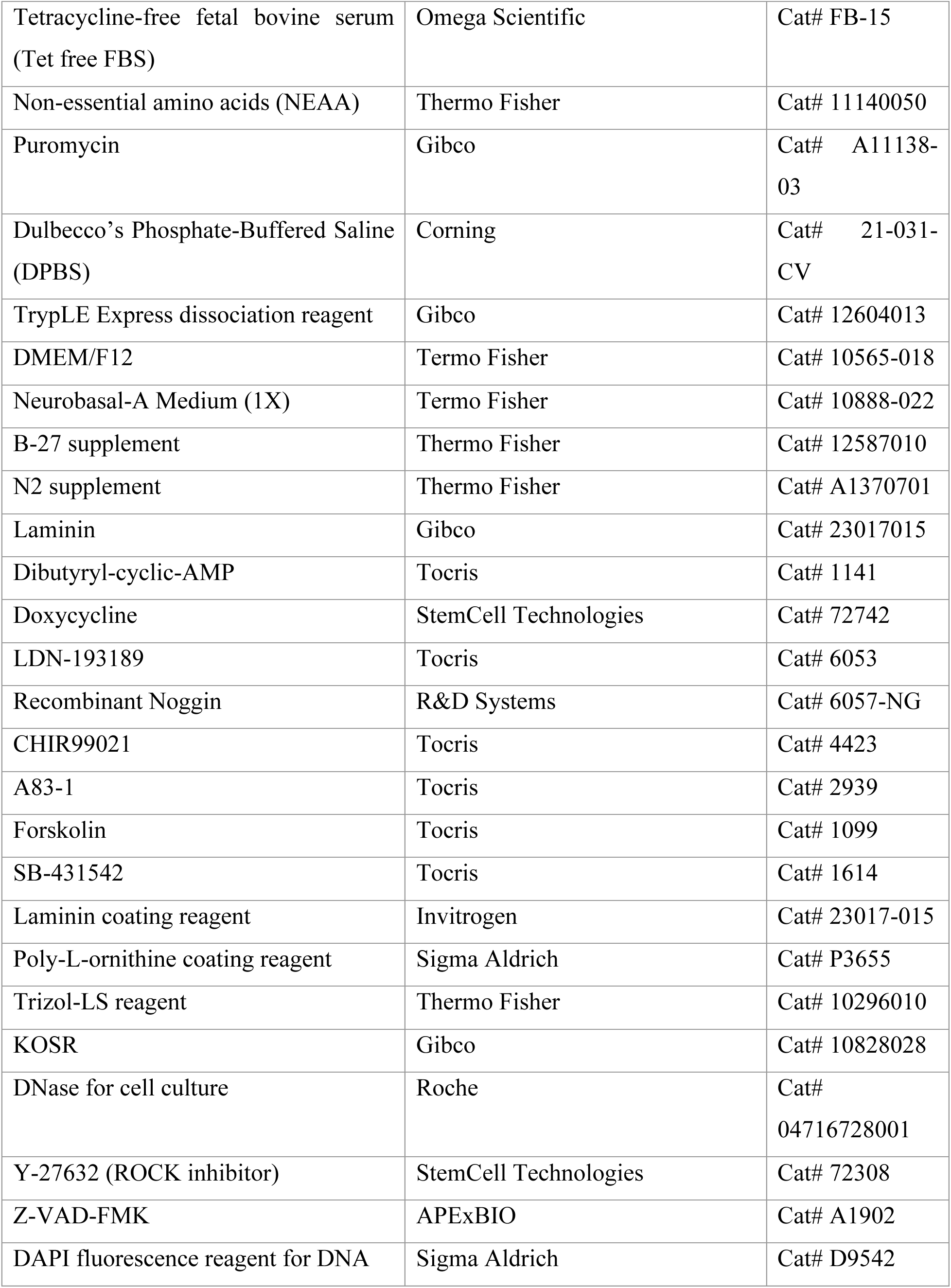
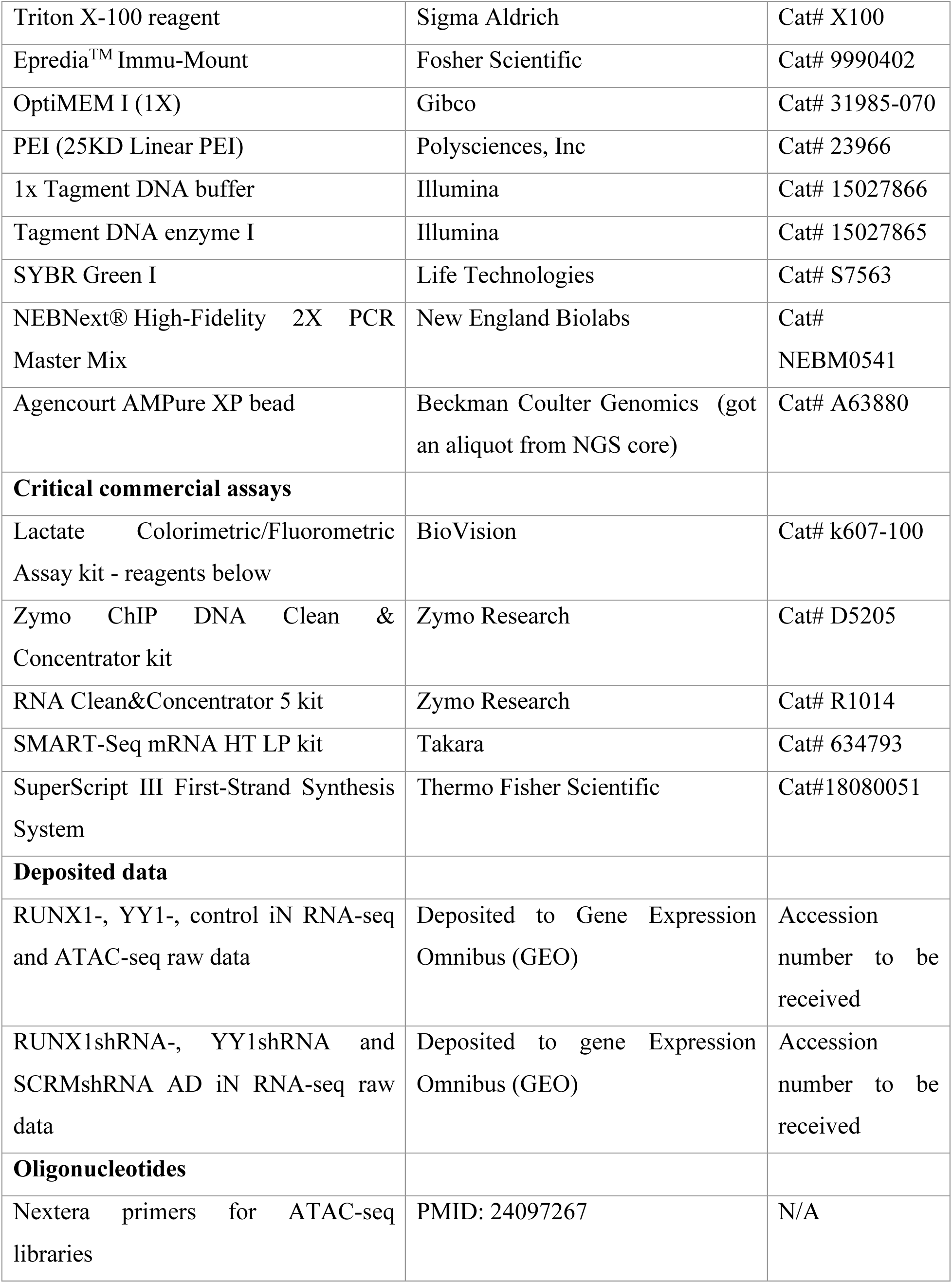
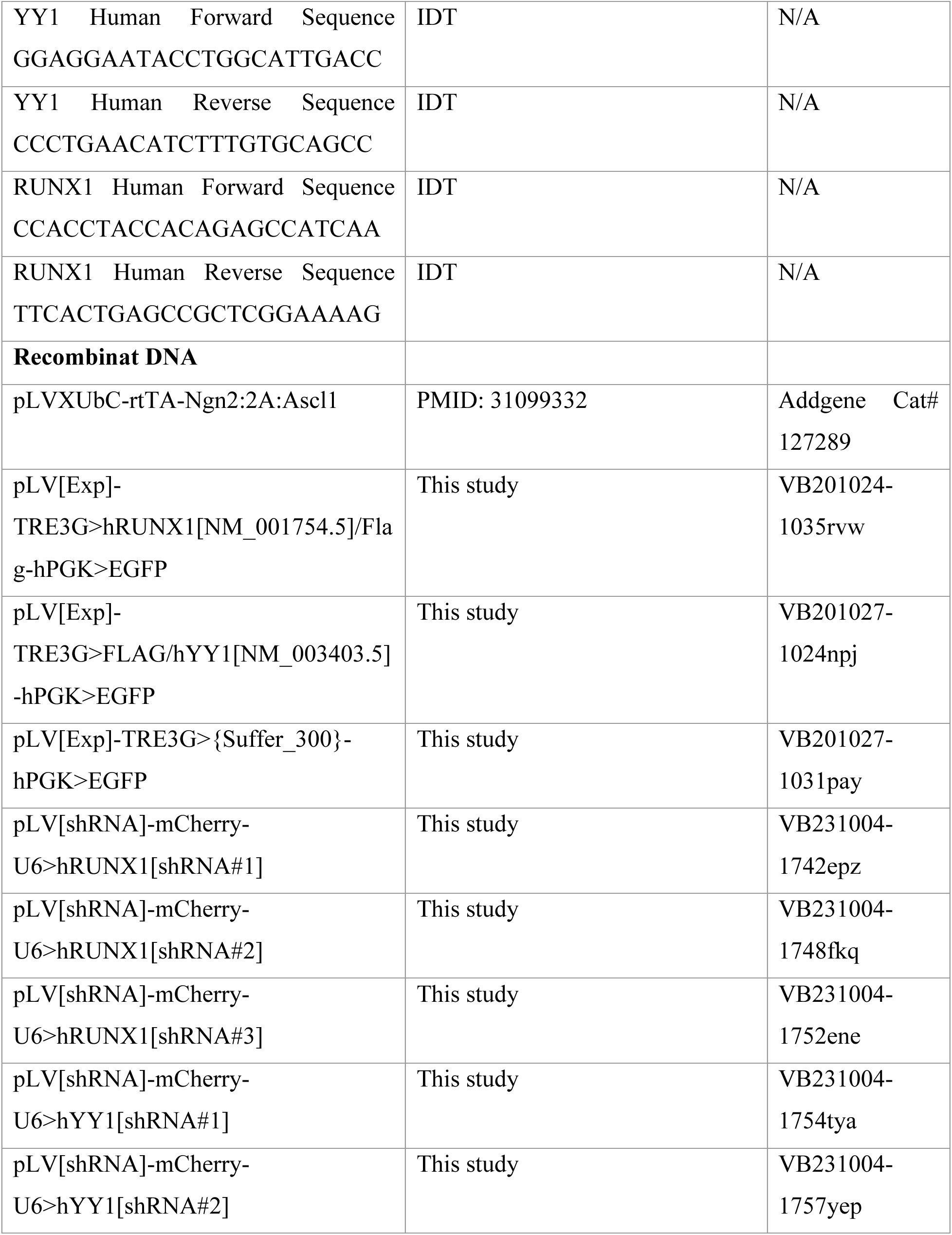
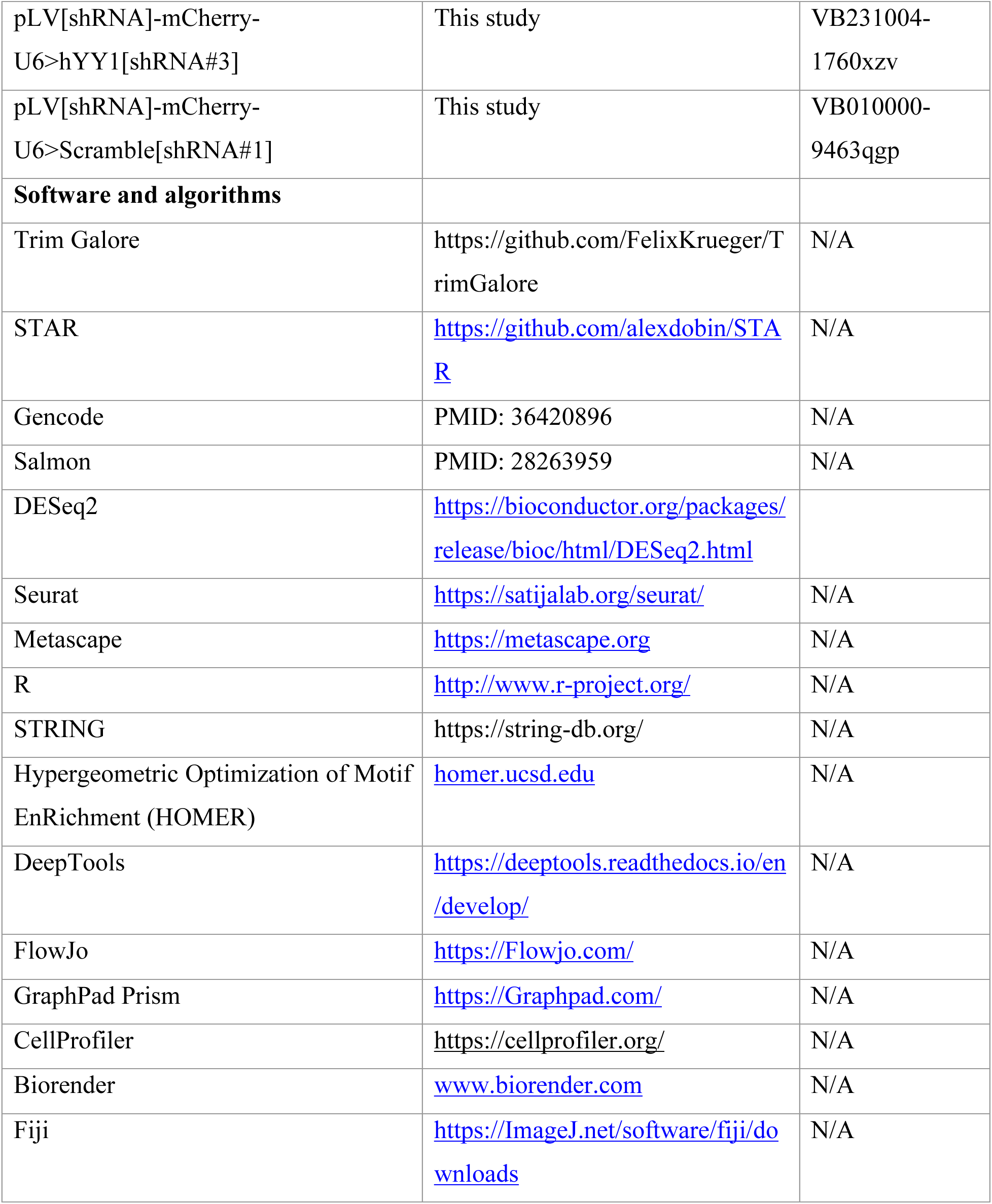

### Materials availability

All unique plasmids and reagents generated in this study are available upon request from the Lead Contact with a complete Materials Transfer Agreement. Lentiviral plasmids for TF OE and KD have been designed and purchased at VectorBuilder, as follow: pLV[Exp]-TRE3G>{Suffer_300}-hPGK>EGFP, pLV[Exp]-TRE3G>hRUNX1[NM_001754.5]/Flag-hPGK>EGFP and pLV[Exp]-TRE3G>FLAG/hYY1[NM_003403.5]-hPGK>EGFP were used for TF OE; pLV[shRNA]-mCherry-U6>Scramble[shRNA]; pLV[shRNA]-mCherry-U6>hRUNX1[shRNA#1]; pLV[shRNA]-mCherry-U6>hRUNX1[shRNA#2]; pLV[shRNA]-mCherry-U6>hRUNX1[shRNA#3]; pLV[shRNA]-mCherry-U6>hYY1[shRNA#1]; pLV[shRNA]-mCherry-U6>hYY1[shRNA#2] and pLV[shRNA]-mCherry-U6>hYY1[shRNA#3] were used for TF KD.

### Data and code availability

All RNA-Seq and ATAC-Seq raw data are being deposited to NCBI Gene Expression Omnibus (GEO) and accession number will be available upon paper acceptance. This paper does not report original code. Any additional information required to reanalyze the data reported in this paper is available from the lead contact upon request.

## EXPERIMENTAL MODEL AND SUBJECT DETAILS

### Subjects

Donors were participants at the Shiley-Marcos Alzheimer’s disease research center (ADRC) at UCSD, who provided a written informed consent. All procedures were approved by the Salk Institute Institutional Review Board. The subjects in this study were obtained from a well characterized cohort of AD and age- and sex-matched non-demented individuals subjected to rigorous clinical assessments, which included a detailed neuropsychological testing and brain magnetic resonance imaging (MRI), as part of the ADRC study, and followed for at least two years after biopsy to confirm clinical diagnosis. Biopsies were punch skin biopsies and were obtained at the UCSD Altman Clinical & Translational Research Institute (ACTRI). Fibroblasts were derived from the biopsies following standard procedures in serum-containing media. Supplementary Table 1 contains subject information. Clinical data and post-mortem pathological evaluations of the individuals from the ROSMAP, where all the subjects are brain tissue sample donors, are in Mathys et. al 2023 ^2^.

### Fibroblasts and iNs

Using a previously described TF-based direct cell conversion method, iNs were directly transduced from primary human dermal fibroblasts (PMID: 33910058). Briefly, fibroblasts were cultured in DMEM (Thermo Fisher, #10569-010) containing 15% tetracycline-free fetal bovine serum (Omega Scientific, #FB-15) and 0.1% non-essential amino acids (NEAA) (Thermo Fisher, #11140050). They were transduced with lentiviral particles for pLVXUbC-rtTA-Ngn2:2A:Ascl1 (UNA) and expanded in the presence of puromycin (Sigma Aldrich, 1μg/ml) as ‘iN-ready’ fibroblast cell lines. Following at least three passages after viral transduction, ‘iN-ready’ fibroblasts were pooled into high densities (approximately 3:1 split from a confluent culture) and, after 24 hours, the medium was changed to neuron conversion medium (NC) based on DMEM:F12/Neurobasal (1:1) for three weeks. NC medium was supplemented with N2 supplement (1x, Thermo Fisher), B27 supplement (1x, Thermo Fisher), doxycycline (2 μg/ml, Sigma Aldrich), laminin (1 μg/ml, Thermo Fisher), dibutyryl-cyclic-AMP (400 μg/ml, Sigma Aldrich), human recombinant Noggin (150 ng/ml, Preprotech), LDN-193189 (500 nM, Fisher Scientific Co), and A83-01 (500 nM, Santa Cruz Biotechnology Inc), CHIR99021 (3 μM, LC Laboratories), Forskolin (5 μM, LC Laboratories) and SB-431542 (10 μM, Cayman Chemicals). After ∼ three weeks of conversion, cells were transduced using lentiviral particles containing a specifically designed plasmid to either overexpress or KD YY1. Twenty-four hours after transduction, the media was then changed to fresh neuronal conversion media and was changed every other day for ∼ one week. Cells were then detached with TrypLE^TM^ Express (Gibco) and Fluorescence-Activated Cell Sorting (FACS)-sorted or plated on tissue culture-treated, poly-L-ornithine (10 μ g/10 mL; Sigma) and laminin (5 μg/mL; Invitrogen) coated ibidi μ-plates (Ibidi) for downstream readouts.

### Method details

#### Multi-omics data integration

We obtained the results from the DE analysis between AD and healthy aged donors in excitatory neurons performed by Mathys and colleagues using the ROSMAP cohort (PMID: 37774677). Next, we narrowed down the list of upregulated genes in AD by selecting only genes with known transcription factor activity (N=1,639 TFs) (PMID: 29425488) and identified 444 upregulated TFs in AD excitatory neurons (adjusted p-value < 0.05 and positive log2 fold-change). In parallel, we also observed that 53 TFs are upregulated in AD iNs (Mertens et al 2021), in which 22 TFs are upregulated in both cohorts of AD patients. To evaluate the gene regulatory network of these TFs, we used DoRothEA ^122^ to retrieve the list of target genes for each of these 22 TFs and performed a Gene-Set Enrichment Analysis (GSEA) on the ROSMAP cohort of excitatory neurons. Subsequently, we selected the TFs with adjusted p-value < 0.05 and positive normalized enrichment score (NES) and identified six candidates. Open chromatin states in AD iNs identified by ATAC-seq (Mertens et al 2021) showed the enrichment of two out of six TFs in open promoter regions associated with AD, as found by a DNA motif search performed using HOMER (adjusted p-value < 0.05).

#### Single cell RNA-sequencing (scRNA-seq) data analysis

DE analysis between AD and healthy aged individuals from the ROSMAP cohort was performed by Mathys and colleagues (PMID: 37774677). We selected the results for excitatory neurons from the group-based differential expression analysis, which compared samples from Group 4 (Pathologic diagnosis of AD and Cognitive diagnosis of Alzheimer’s dementia) versus Group 1 (No pathologic diagnosis of AD and No cognitive impairment). We used the adjusted *p*-value and log2 fold-change provided by the authors as part of the supplemental material of the manuscript.

#### Plasmid construct design

The following plasmids were designed for TF OE in healthy aged iNs using VectorBuilder: RUNX1 plasmid pLV[Exp]-TRE3G>hRUNX1[NM_001754.5]/Flag-hPGK>EGFP, YY1 plasmid pLV[Exp]-TRE3G>FLAG/hYY1[NM_003403.5]-hPGK>EGFP and control plasmid pLV[Exp]-TRE3G>{Suffer_300}-hPGK>EGFP (Supplementary Figure 2A).

Using VectorBuilder, the following plasmids were then designed for TF KD in AD iNs: three separate RUNX1shRNAs, such as pLV[shRNA]-mCherry-U6>hRUNX1[shRNA#1], pLV[shRNA]-mCherry-U6>hRUNX1[shRNA#2] and pLV[shRNA]-mCherry-U6>hRUNX1[shRNA#3], three separate YY1shRNAs, such as pLV[shRNA]-mCherry-U6>hYY1[shRNA#1], pLV[shRNA]-mCherry-U6>hYY1[shRNA#2] and pLV[shRNA]-mCherry-U6>hYY1[shRNA#3] and one scramble (SCRM) shRNA, such as pLV[shRNA]-mCherry-U6>Scramble[shRNA] (Supplementary Figure 6A). After being tested for transduction efficiency using Human Embryonic Kidney 293 (HEK293) cells, RUNX1shRNA plasmid pLV[shRNA]-mCherry-U6>hRUNX1[shRNA#1] and YY1 shRNA plasmid pLV[shRNA]-mCherry-U6>hYY1[shRNA#1] were selected out of three RUNX1 and YY1 shRNAs.

All plasmids were incorporated into lentiviral particles by the Salk Gene Transfer, Targeting and Therapeutics Viral Vector Core and used for either permanently OE or KD RUNX1 or YY1.

#### Permanent overexpression

iNs generated from heathy aged subjects were transduced with lentiviral particles containing either RUNX1 plasmid pLV[Exp]-TRE3G>hRUNX1[NM_001754.5]/Flag-hPGK>EGFP, YY1 plasmid pLV[Exp]-TRE3G>FLAG/hYY1[NM_003403.5]-hPGK>EGFP or control plasmid pLV[Exp]-TRE3G>{Suffer_300}-hPGK>EGFP at ∼ day 19 of iN conversion. Media was changed to fresh NC media following 24h of transduction and iNs were cultured in neuronal conversion media for ∼ eight days before fixing and staining, or isolation by FACS for downstream readouts.

#### Permanent knockdown

iNs directly generated from AD subjects were transduced with lentiviral particles containing either RUNX1shRNA plasmid pLV[shRNA]-mCherry-U6>hRUNX1[shRNA#1]), YY1 shRNA plasmid pLV[shRNA]-mCherry-U6>hYY1[shRNA#1] or SCRM shRNA plasmid pLV[shRNA]-mCherry-U6>Scramble[shRNA]. Media was changed to fresh NC media following 24h of transduction and iNs were cultured in neuronal conversion media for ∼ 8 days. Converted iNs expressing the mCherry fluorescent marker were then isolated by FACS for RNA purification and subsequent RNA-seq.

#### Flow cytometry

Following ∼three week conversion and ∼one week after transduction, cells were detached using TrypLE^TM^ Express (Gibco) and stained for PSA-NCAM directly conjugated to APC (Miltenyi Biotec, 1x) for 1h at 4°C in sorting buffer (250 mM myo-inositol and 5 mg/ml polyvinyl alcohol in PBS) containing 5% KOSR (Thermo Fisher). Cells were washed and resuspended in sorting buffer containing EDTA and DNase and filtered using a 40-μm cell strainer. For the TF gain-of-function study, we sorted for EGFP+/PSA-NCAM+ double-positive cells to select for iNs successfully transduced with lentiviral particles. Specifically, live cells were selected first by gating for DAPI-negative cells, followed by exclusion of debris using forward and side scatter pulse area parameters (FSC-A and SSC-A) and exclusion of aggregates using forward and side scatter pulse width (FSC-W and SSC-W), before gating populations based on PSA-NCAM then EGFP fluorescence. As neuronal cells express the PSA-NCAM marker, cells were first selected for PSA-NCAM. A BD Influx sorter (BD Bioscience) was used to sort cells, with 1x PBS for sheath fluid, using a ’1-drop pure’ mode. A 100-μm nozzle was used for cells with sheath pressure set to 20PSI, with all samples and collection tubes kept at 4°C for RNA-seq or room temperature for replating. PSA-NCAM was excited with a 640nm laser (120mW for Influx) and detected using the “APC” channel (670/30BP). EGFP was excited with a 488nm laser (200mW) and detected using the “FITC” channel (530/40BP). For TF loss-of-function study, we sorted for mCherry+/PSA-NCAM+ double-positive cells instead. PSA-NCAM+ cells were sorted as described above and subsequently selected for mCherry instead of EGFP. The latter was excited with a 561nm laser (150mW for Influx) and detected using the “PE” channel (593/40BP).

For RNA isolation, cells were sorted into Trizol-LS reagent (Thermo Fischer). For ATAC-seq, cells were sorted into PBS (Corning). For extended culture, iNs were sorted into NC media containing DNase, Rock-inhibitor (10 *μ*M), z-VAD-FMK (20 *μ*M) and replated at 150 cells/cm2 on poly-L-ornithine/laminin coated ibidi 96-well plates.

#### RNA isolation

Total bulk RNA was extracted from FACS-sorted iNs expressing EGFP or mCherry, following three weeks of conversion and one week after transduction, using Trizol-LS reagent (Thermo Fischer) and a chloroform phase separation method coupled with the RNA Clean&Concentrator 5 RNA extraction kit (Zymo Research), according to the manufacturer’s instructions. Before RNA-seq library preparation, RNA integrity was assessed using RIN Agilent TapeStation HighSensitivity RNA Screen Tape (Agilent) or 2100 Bioanalyzer Instrument (Novogene).

#### mRNA-sequencing (RNA-seq) and data analysis

cDNA libraries were generated using the SMART-Seq mRNA HT LP Sample Prep Kit according to the manufacturer’s instructions (Takara). One hundred bp paired-end (PE) sequencing was performed using the Illumina NovaSeq6000 platform or the Illumina NovaSeq X Plus 1.5B Flowcell platform. Three replicates per line for three healthy aged lines overexpressing RUNX1, two healthy aged lines overexpressing YY1 and three control healthy aged lines were subsequently sequenced. Low-quality (Phred score < 20) ends from reads were trimmed using Trim Galore (version 0.6.7) (https://github.com/FelixKrueger/TrimGalore) in addition to adapter removal. Reads were then aligned to the human reference genome (GRCh38.p13) using STAR (version 2.7.11b) (PMID: 23104886) and annotated using the Gencode gene annotation (version 43) (PMID: 36420896). Counts and Transcripts Per Million (TPM) values were quantified using Salmon (version 1.9.0) (PMID: 28263959) at gene level (N=19,084 features). Differential gene expression analysis between iNs OE each TF and control was performed using the Bioconductor package DESeq2 (version 1.46, PMID: 25516281), incorporating the replicate as a covariate. All statistical tests were followed by multiple testing corrections using the Benjamini and Hochberg (BH) method for False Dicovery Rate (FDR) estimation. Differentially expressed genes were identified using an adjusted *p*-value threshold of 0.01 and log2 Fold-change (FC) > 1 for upregulated genes, and log2 FC < -1 for downregulated genes. For the loss-of-function study, shRNA expression cassettes were used to KD YY1 and RUNX1, separately, in independent iN cultures generated from three Alzheimer’s disease (AD) lines, along with paired control samples treated with SCRM shRNA. Samples were processed as described above. DE genes were identified using an adjusted *p*-value threshold of 0.01 and log2 FC > 0 for upregulated genes and log2 FC < 0 for downregulated genes.

#### GO analysis and gene network analysis

Functional annotation for GO biological processes, cellular components, molecular function and KEGG pathways was performed using Metascape (PMID: 30944313). In Metascape-based analysis, the reported *P* values are calculated using the hypergeometric test. The statistical significance threshold for all GO analyses was *P* < 0.01. Gene interaction network analysis was performed using STRING. In these plots, each point is a gene, or node, and gene-gene connections, or edges, are shown as colored lines, indicating the type of interaction evidence as previously described (string-db.org).

#### Assay for Transposase-Accessible Chromatin with high-throughput Sequencing (ATAC-seq) and data analysis

ATAC-Seq was performed as previously described ^43,44^. Briefly, 50,000 FACS-purified iNs generated from four independent aged healthy lines were lysed in 50 μl lysis buffer (10 mM Tris-HCl ph 7.5, 10 mM NaCl, 3 mM MgCl2, 0.1% IGEPAL, CA-630, in water), on ice and pelleted in a fixed-angle refrigerated benchtop centrifuge. Collected nuclei were then resuspended in 50 μL transposase reaction mix (1x Tagment DNA buffer, 2.5 μL Tagment DNA enzyme I in water (Illumina), and incubated at 37°C for 30 min. DNA was purified with a Zymo ChIP DNA Clean & Concentrator Kit (Zymo Research D5205) according to the manufacturer’s instructions. DNA was then amplified with PCR mix (1.25 μM Nextera primer 1, 1.25 μM Nextera primer 2-bar code, 0.6x SYBR Green I (Life Technologies, S7563), 1x NEBNext High-Fidelity 2x PCR MasterMix, (NEBM0541) for 7-10 cycles. Amplified libraries were purified using AMPure XP beads (Beckman Coulter Genomics, A63880) for fragment size selection according to the Illumina Nextera Kit recommended protocol. Libraries were quantified using Qubit HS DNA Assay on a Qubit 3.0 Fluorometer (Thermo Fisher Scientific) according to the manufacturer’s instructions. Library sizes were assessed on an Agilent Tape Station High Sensitivity DNA Screen Tape using the High Sensitivity DNA chip (Agilent Technologies). PE100 bp sequencing was performed using the Illumina NovaSeq6000 platform. Reads were trimmed using TrimGalore (https://github.com/FelixKrueger/TrimGalore) and mapped to the UCSC genome build hg38 using STAR. Peaks were then calculated and annotated using the HOMER software package ^123^. For agnostic genome-wide characterization of the peaks, sequence depth-normalized bigWig files were used for generating chromatin accessibility profiles using deepTools software ^124^. Next, differential accessibility analysis of all identified ATAC-Seq peaks was performed using differential peaks function in HOMER, and fold changes and significance of the resulting differential peaks were plotted as a volcano plot using R ggplot2. Genome-wide integration of the ATAC-Seq data with the RNA-Seq data on a gene-by-gene level was performed based on comparing the fold changes of differentially accessible peaks (HOMER differential ATAC peaks), with the fold changes of DE genes (RNA-Seq expression), using R, and visualization was performed using R ggplot2. Motif enrichment analysis was performed in HOMER. The biological relevance of TFs with high binding affinity for annotated Promoter-TSS peaks was assessed using Metascape.

#### Immunocytochemistry

Cells were fixed with 4% paraformaldehyde for 30 min at room temperature and washed 3 x 15 min with TBS, followed by a 30 min block with TBS containing 3% serum and 0.1% Triton X-100. Cells were incubated with primary antibodies NeuN (1:500; EMD Millipore), β-III-tubulin (1:500; Biolegend), p-PKM2(Ser37) (1:500; SAB Signalway Antibody), EGFP (1:2000; Aves Lab) at 4°C overnight. Following two 5-minute washes with TBS and 30 minutes in TBS containing 10% serum and 0.1% Triton X-100, cells were incubated with secondary antibodies (Donkey Anti-Chicken IgG Alexa Fluor 647, Cy3 Donkey Anti-Mouse IgG, Cy3 Donkey Anti-Rabbit IgG, Donkey Anti-Mouse IgG Alexa Fluor 647 and Donkey Anti-Chicken IgG Alexa Fluor 488, all 1:500) for 2 hours at room temperature. After washing, nuclei were stained with DAPI (1:10,000; Sigma-Aldrich). Following one wash in TBS containing 10% serum and 0.1% Triton X-100 and one wash in TBS, plates were mounted in Epredia^TM^ Immu-Mount (Fisher Scientific). Confocal images were acquired on standard fluorescence microscopes. Expression of pPKM2 was assessed in two replated FACSorted iN lines OE YY1 including three to five technical replicates each, and four control iN lines including one to eight technical replicates each. pPKM2 nuclear expression was measured as Integrated Density (IntDen) in regions of interests (ROIs) set based on DAPI and total neuronal expression was measured as IntDen in ROIs based on TUJ1. All data for one experiment were acquired from cells cultured and processed in parallel on the same microscope with the exact same settings used.

#### Neuronal morphometric reconstruction and analysis

Automated Sholl analysis was performed using the open-source image analysis software Cellprofiler (PMID: 17269487). Nuclei were identified based on minimum cross-entropy global thresholding of DAPI intensity using the “IdentifyPrimaryObjects” module. Nuclear objects were expanded by a 25-pixel (approximately 5-micron) radial distance to define the soma. The immunofluorescence intensity of the neuronal marker TUJ1 was measured within soma and TUJ1-positive soma were filtered and designated as neuronal soma. Neurites in the TUJ1 immunostain were enhanced using the “EnhanceOrSuppressFeatures” module. The “IdentifySecondaryObjects” module was used to identify total neuronal cell area by propagating the previously identified neuronal soma along the enhanced neurite image using two-class Otsu adaptive thresholding. The enhanced neurite image was binarized and skeletonized to generate a linear neurite network. Total neurite network length was measured using the “MeasureObjectSizeShape” module and measurements were related to previously defined individual neuronal cell areas using the “RelateObjects” module. The Sholl radial grid was built by expanding outward from the nuclear objects a series of ten 25-pixel-thick annuli, corresponding to Sholl radii. Skeletonized neurite objects within each annulus were masked to define object intersections at known Sholl distances. Since neurite objects from different cells can cross through Sholl radii of adjacent cells, neurite objects within each annulus were related back to individual neuronal cell areas using the “RelateObjects” module to disambiguate single-cell measurements. Data are presented as the mean and standard error of >150 cells/condition from 3 independent experiments and unique donors.

#### Lactate Colorimetric Assay

Following three weeks of neuronal conversion and one week of transduction, FACS-sorted YY1-and control iNs were plated on poly-L-ornithine (10 μ g/10 mL; Sigma) and laminin (5 μg/mL; Invitrogen)-coated black-walled ibid 96-well plates (Ibidi). Three days after re-plating, supernatant was collected and analyzed using colorimetric lactate assay according to the manufacturers’ instructions (BioVision). Recording of signals was performed using the GloMax Discover Plate Reader platform (Promega).

#### Quantification and statistical analysis

Data analysis and data visualization were performed using R (version 4) software packages (www.r-project.org) and Bioconductor. Unless specified, all statistical tests were performed using two-tailed tests and significance was obtained with FDR < 5%. GraphPad Prism software was used to calculate non-omics quantitative data statistics using the method indicated in each figure or method section. Statistical significance is marked as \**p* < 0.05; \*\**p* < 0.01; \*\*\**p* < 0.001; \*\*\*\**p* < 0.0001 in the figures.

## Acknowledgments

We are grateful to all donors participating in this study. We further thank Mary Lynn Gage for editorial comments. Illustrations were created using BioRender.com. This work was supported by the National Institute on Aging (NIA) 4 RF1 AG056306-07; California Institute for Regenerative Medicine (CIRM) grant INFR6.2-15440; the Milky Way Research Foundation; the Ray and Dagmar Dolby Family Fund; Carl C. Anderson Sr & Marie Jo Anderson Foundation; Salk Women & Science Scientific Career and Professional Development award; the AHA-Allen Initiative in Brain Health and Cognitive Impairment award made jointly through the American Heart Association and The Paul G. Allen Frontiers Group: 19PABH134610000; NIH R01 AG056511; BrightFocus Foundation award A2022024F; Live like Lou Foundation PostDoctoral fellowship; and the Shiley-Marcos Alzheimer’s Disease Research Center (ADRC; AG062429) at the University of California, San Diego (UCSD); the Flow Cytometry Core Facility of the Salk Institute (RRID:SCR_014839) with funding from NIH-NCI CCSG P30 CA014195, and Shared Instrumentation Grants S10-OD023689 (Aria Fusion cell sorter), and S10 OD034268 (Thermo Fisher Bigfoot); the NGS Core Facility of the Salk Institute with funding from NIH-NCI CCSG: P30 014195; the Waitt Advanced Biophotonics Core Facility of the Salk Institute (RRID:SCR_014838) with funding from NIH-NCI CCSG P30 CA014195, NIH-NIA San Diego Nathan Shock Center P30 AG068635, The Henry L. Guenther Foundation and the Waitt Foundation; the GT3 Core Facility of the Salk Institute with funding from NIH-NCI CCSG: P30 CA014195; the Stem Cell Core Facility of the Salk Institute.

## Author contributions

Conception, design, and writing of the manuscript: R.L. and F.H.G. Performance, analysis and/or interpretation of cellular and molecular assays: R.L., Y.V., L.K., L.T, A.K., J.J and M.R. Next-generation sequencing and bioinformatics analysis: T.S.S, R.L., M.C and J.R.H. Data interpretation and editing of the manuscript: R.L. and F.H.G, with comments from J.M., J.R.H. and S.T.S.

---

**Supplementary Figure 1.**
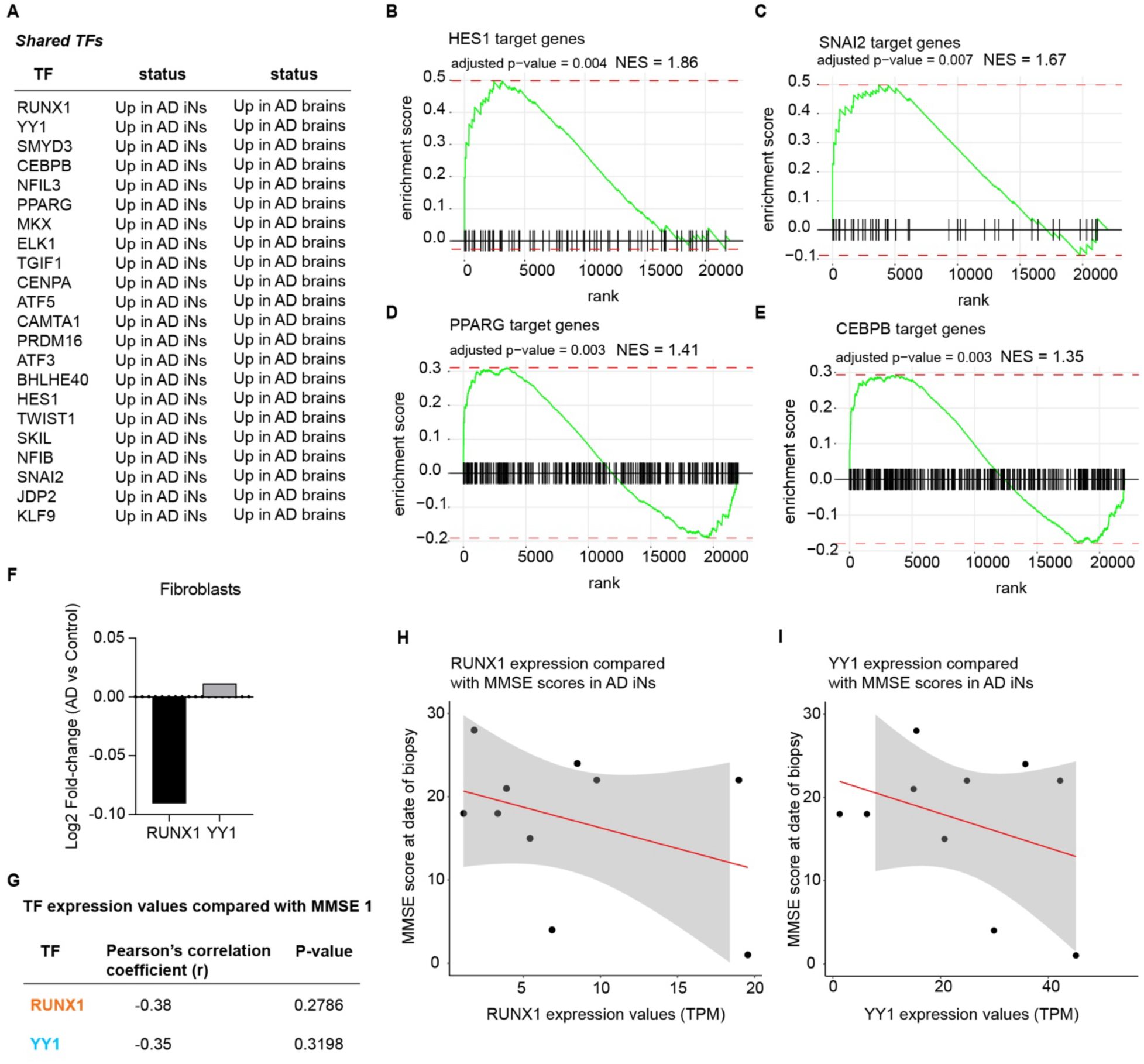
TFs dysregulated in human postmortem AD brains and AD iNs. (A) List of 22 TFs significantly upregulated in both AD iNs and human post-mortem AD brains. (B-E) GSEA of HES1, SNAI2, PPARG and CEBPB target genes in excitatory neurons from human post-mortem AD brains. (F) RUNX1 and YY1 were not differentially significantly expressed in AD fibroblasts from the same donors. (G) Correlation values of RUNX1 and YY1 expression to MMSE and corresponding adjusted *p*-values in AD iNs. (H-I) Pearson correlation between *RUNX1* and *YY1* expression and MMSE of the AD and control individuals was performed to assess the correlation between TF expression and cognitive function.

**Supplementary Figure 2.**
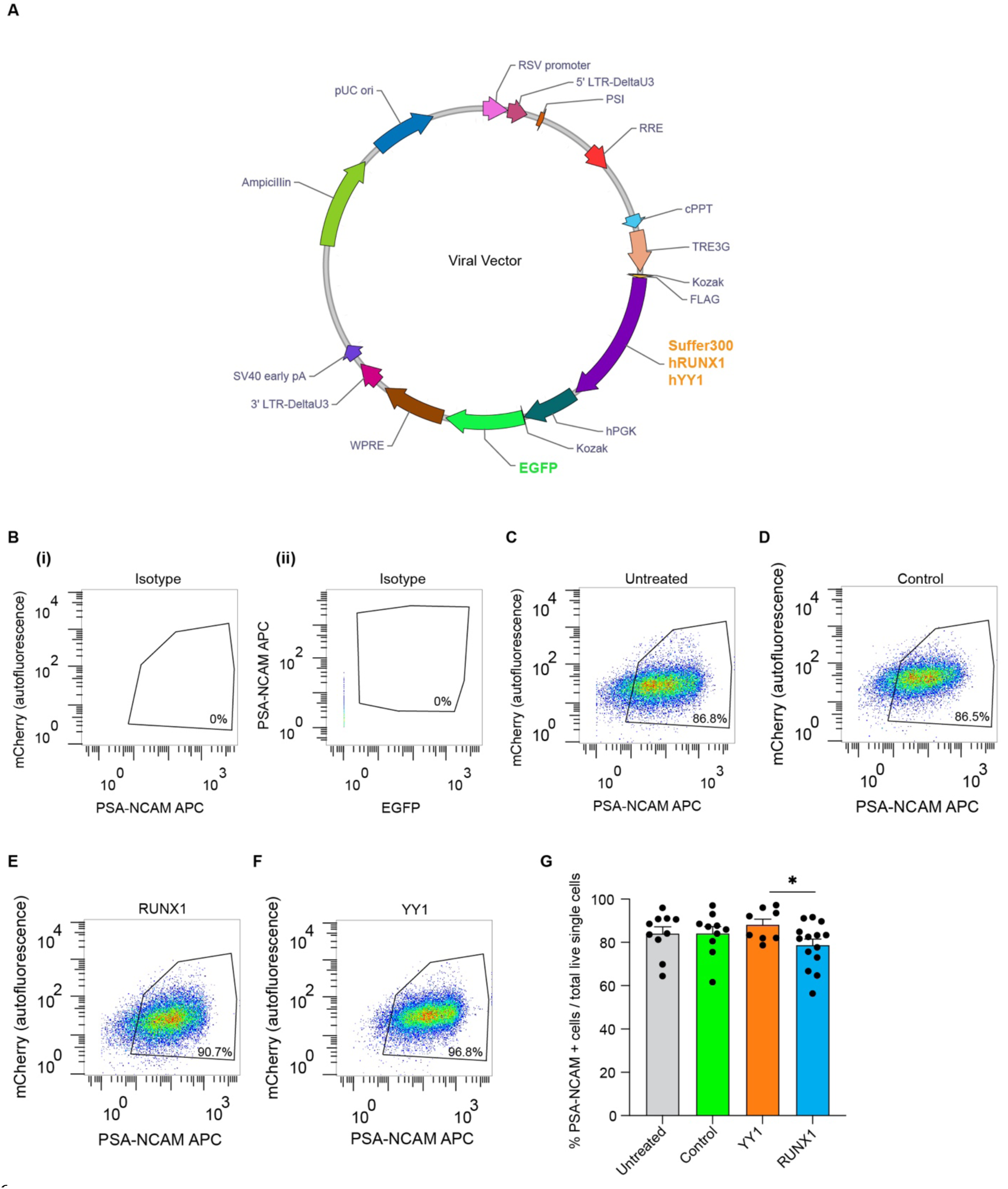
TF overexpression in healthy aged iNs. (A) Plasmid map of the viral vector construct used to overexpress either RUNX1, YY1 or control (Suffer300) sequence. (B-F) Representative FACS gates to illustrate gating strategy. iNs were sorted using neuronal marker PSA-NCAM, using mCherry for removing autofluorescence, and by EGFP. Example plot and gating for FACS-based purification of PSA-NCAM+ cells, using isotype control (B), from untreated iNs (C), iNs OE control sequence (D), iNs OE RUNX1 (E) and iNs OE YY1 (F). (G) FACSorted PSA-NCAM+ cells over live single cells (*n* = 3 x Untreated, *n* = 3 x control, *n* = 3 x iNs OE YY1, *n* = 3 x iNs OE RUNX1 and 1 to 5 independent iN culture replicates per condition). Statistical significance was assessed by performing one-way ANOVA, Tukey’s multiple comparisons test.

**Supplementary Figure 3.**
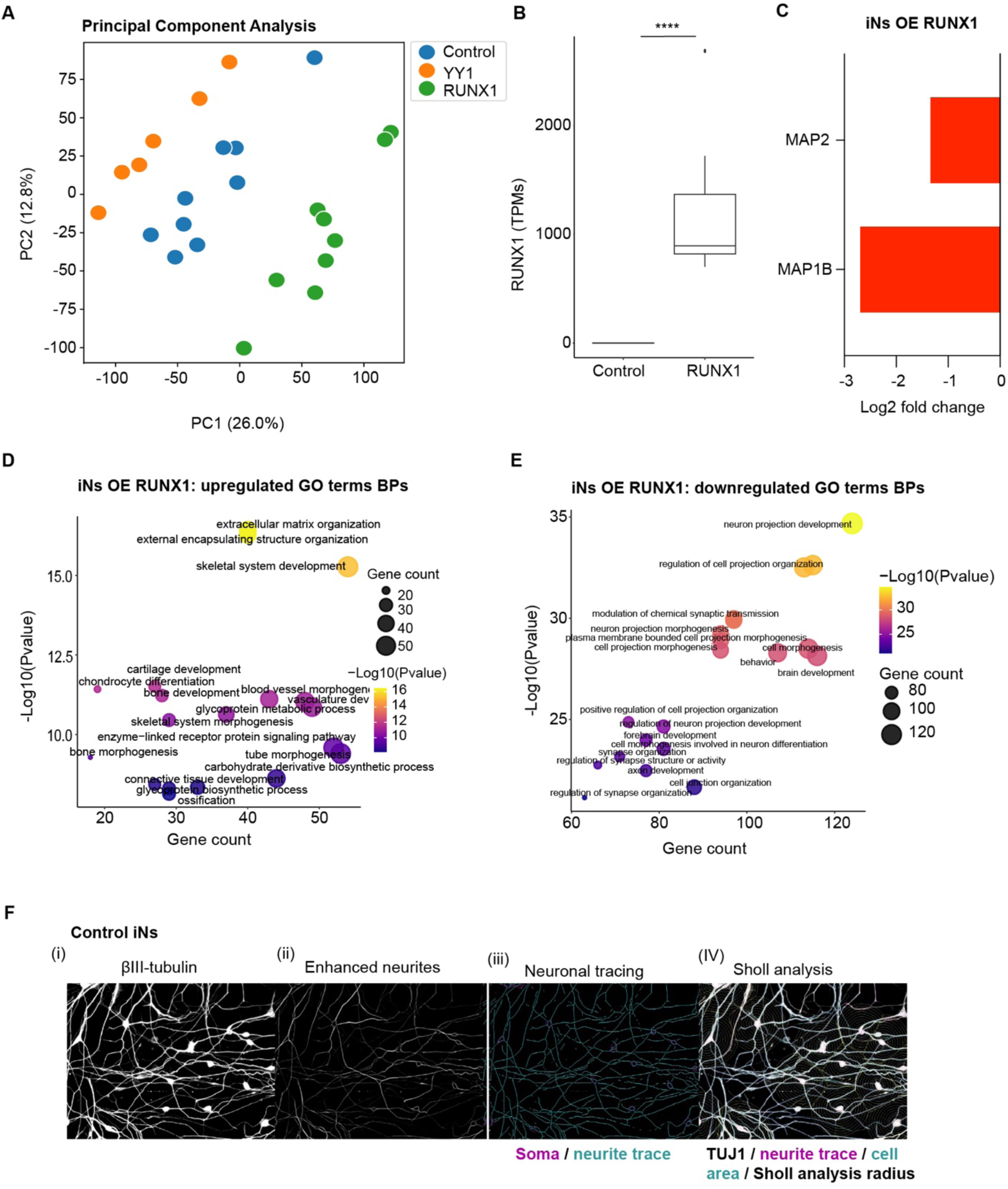
Transcriptional and phenotypical changes driven by RUNX1 OE in aged iNs. (A) Principal component analysis was performed on RNA-seq data from the iNs OE RUNX1 versus control, and YY1 versus control (3 independent iN culture replicates per line for *n* = 3 iNs OE RUNX1, *n* = 2 iNs OE YY1 and *n* = 3 controls). PC1 and PC2 explained the largest variance among the samples and revealed a clear separation between each group. (B) RUNX1 expression values (TPM, transcript per millions) in the iNs OE RUNX1 and control (* *p*adj < 0.05, ** *p*adj < 0.01, *** *p*adj < 0.001, **** *p*adj < 0.0001). (C) MAP2 and MAP1B expression in aged iNs following RUNX1 OE. (D-E) Top significantly upregulated (D) and downregulated GO terms biological processes (BPs) in iNs OE RUNX1. Point sizes correspond to the number of genes assigned to each category (Supplementary Table 2). (F) Neuronal morphometric reconstruction of control iNs. (i) Representative immunofluorescence image of control iNs stained for neuronal marker βIII-tubulin, (ii-iV) neuronal reconstructions and Sholl analysis using Cell Profiler. Scale bar 10 μm. Experiments were performed in all 3 patient lines and 2 iN culture replicates per line.

**Supplementary Figure 4.**
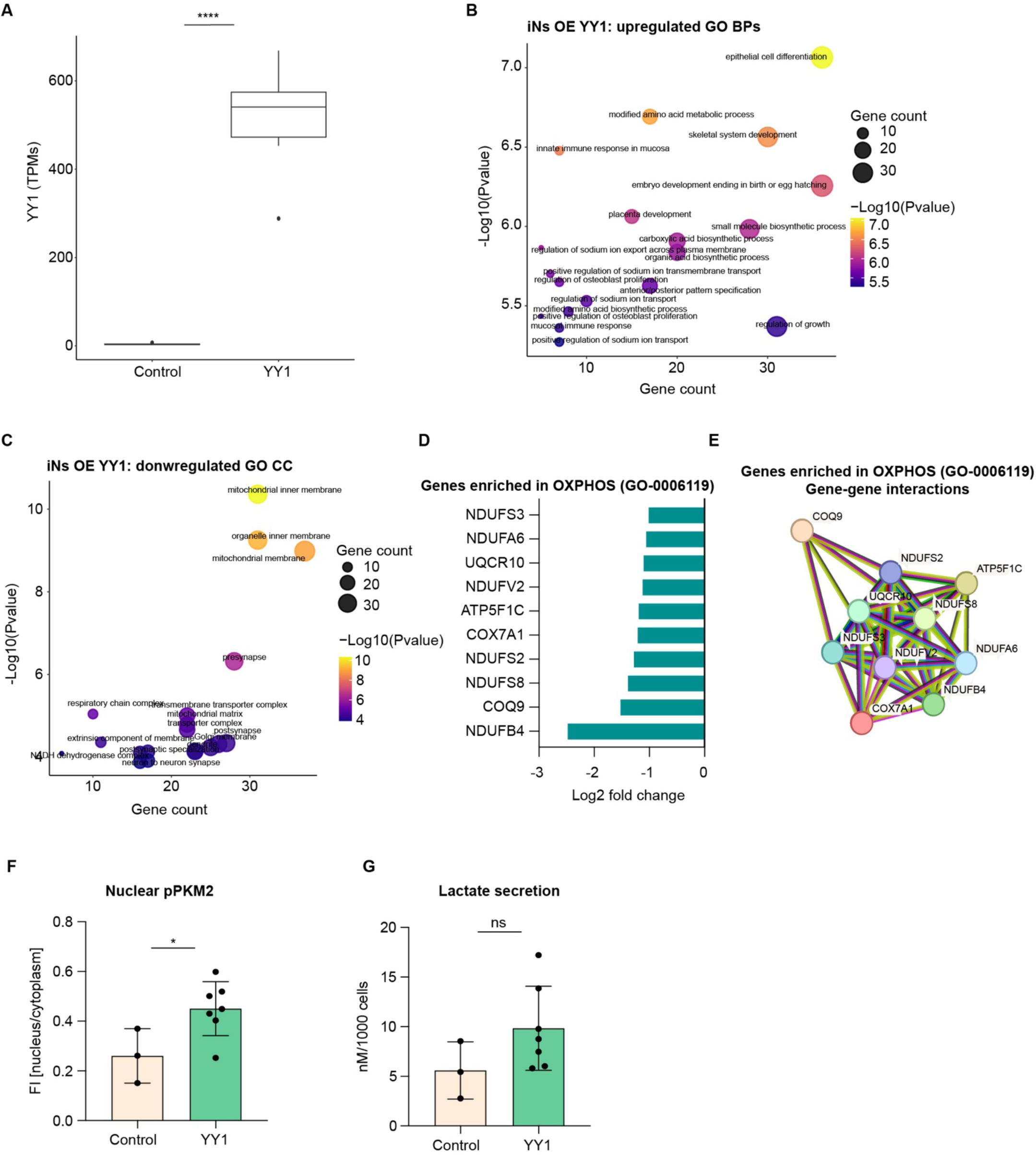
Transcriptional and phenotypical changes driven by YY1 OE in aged iNs. (A) YY1 gene expression values (TPMs) in the iNs OE YY1 and control (* *p*adj < 0.05, ** *p*adj < 0.01, *** *p*adj < 0.001, **** *p*adj < 0.0001). (B) Top significantly enriched GO BPs within significant upregulated genes in iNs OE YY1. Point sizes correspond to the number of genes assigned to each category (Supplementary Table 3). (C) Top significantly downregulated GO CC within significant downregulated genes in iNs OE YY1. Point sizes correspond to the number of genes assigned to each category (Supplementary Table 3). (D-E) Bar graph showing gene expression changes of OXPHOS genes in the iNs following YY1 OE (D) and STRING network showing gene-gene connections across the represented OXPHOS genes (E). (F) Quantification of pPKM2(Ser37) in two independent donors under control condition (EGFP only) and after YY1 OE (each dot represents one independent iN culture; significance, unpaired *t*-test). (G) Colorimetric assay was performed to measure secreted lactate in the supernatant of two independent donors under control condition (EGFP only) and after YY1 OE (each dot represents one independent iN culture; significance, unpaired *t*-test).

**Supplementary Figure 5.**
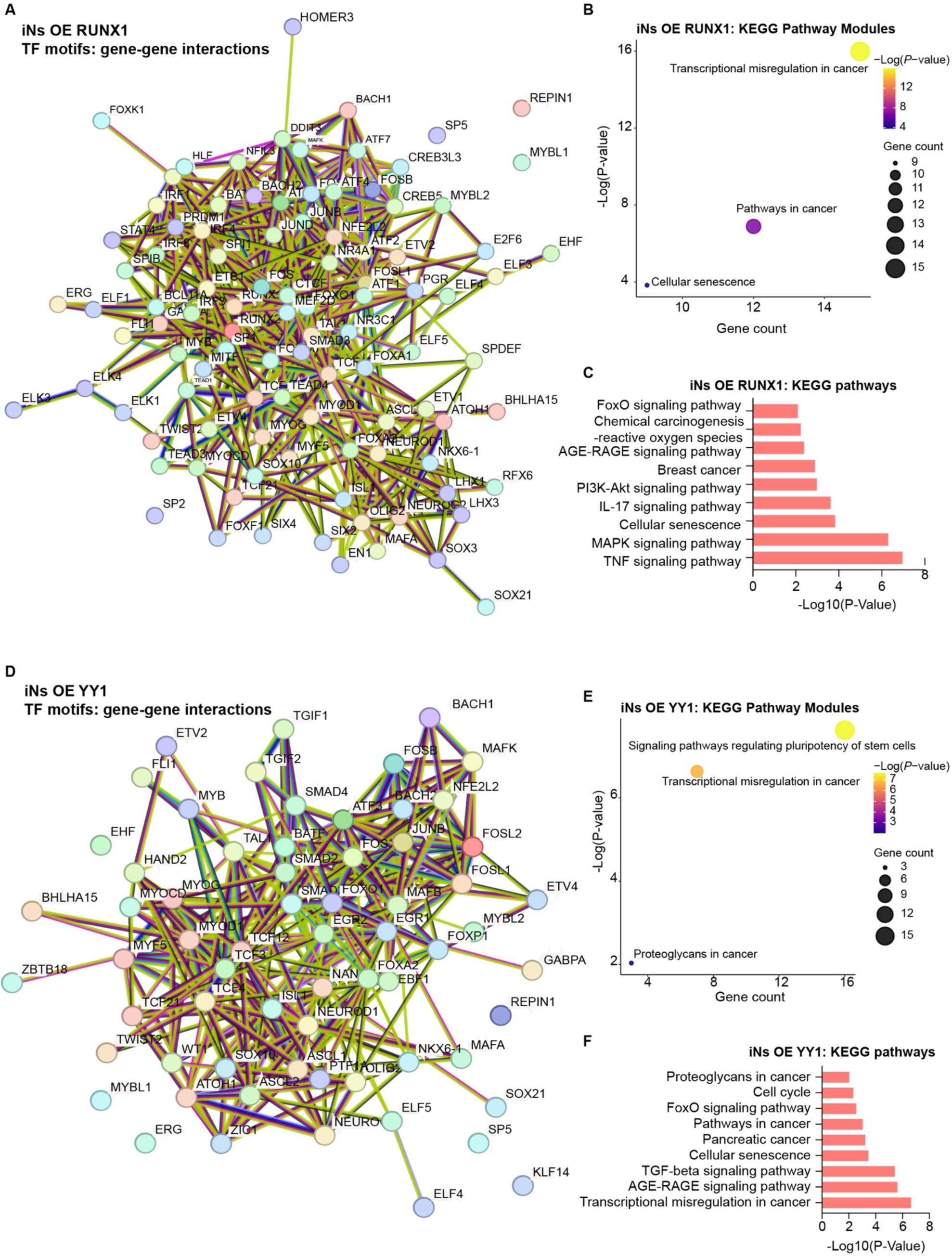
Characterization of the differentially accessible regulatory chromatin landscape following RUNX1 and YY1 OE in aged iNs. (A) STRING network showing gene-gene connections across the TFs with high binding affinity to differentially open *cis*-regulatory sites in the iNs OE RUNX1. (B-C) KEGG pathway analysis of the TFs with high affinity for all significantly enriched motifs in the iNs OE RUNX1. Bubble plots for KEGG pathway modules; point sizes correspond to the number of genes within each cluster, colored by significance (B and Supplementary Table 5). Bar graph shows selected significantly enriched KEGG pathways (C and Supplementary Table 5). (D) STRING network showing gene-gene connections across the TFs with high binding affinity to differentially open *cis*-regulatory sites in the iNs OE YY1. (E-F) KEGG pathway analysis of the TFs with high affinity for all significantly enriched motifs in the iNs OE YY1. Bubble plots for KEGG pathway modules; point sizes correspond to the number of genes within each cluster, colored by significance (E and Supplementary Table 7). Bar graph shows selected significantly enriched KEGG pathways (F and Supplementary Table 7).

**Supplementary Figure 6.**
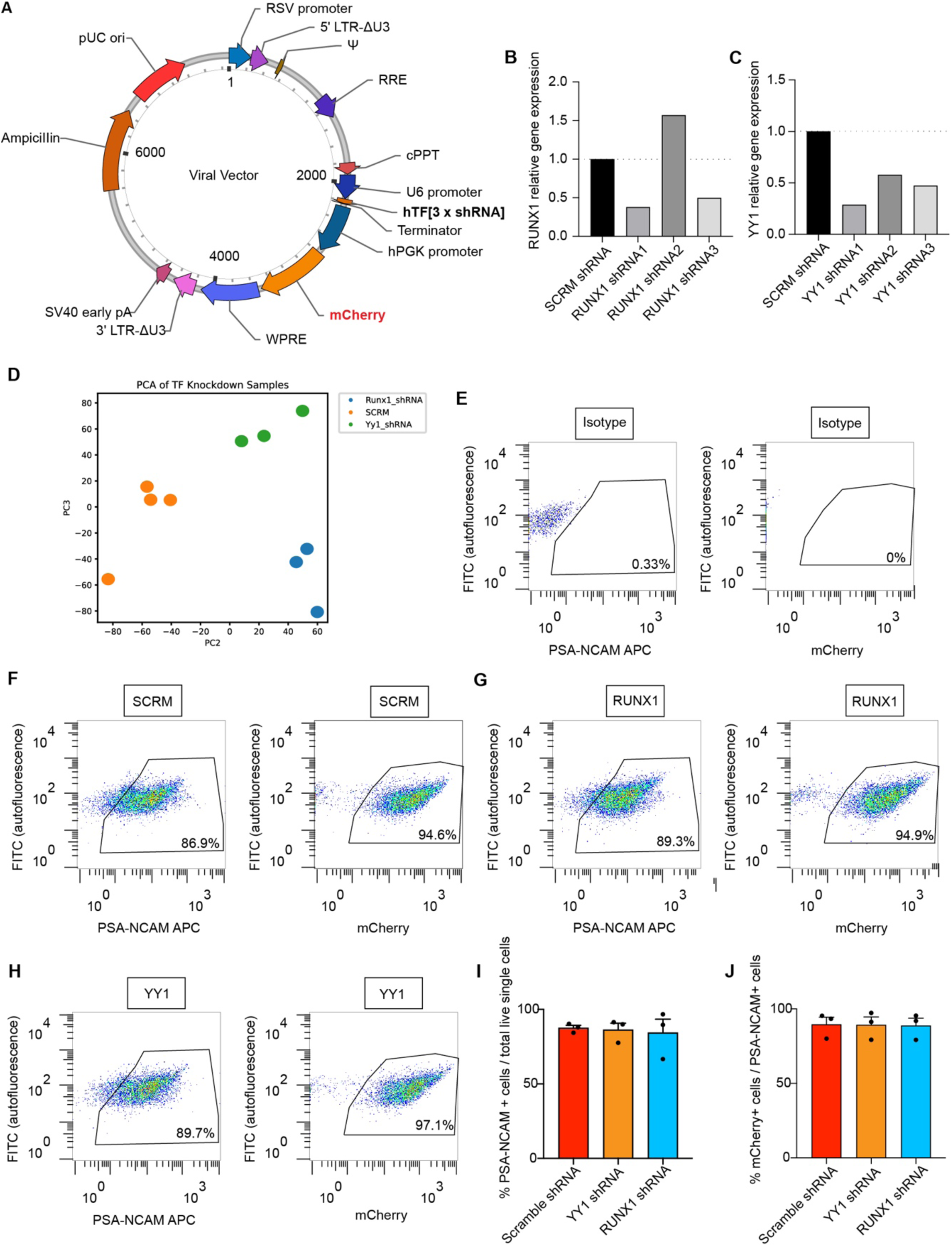
RUNX1 and YY1 downregulation in AD iNs. (A) Plasmid map of the viral vector construct used to downregulate RUNX1 or YY1 expression in AD iNs. (B) Changes in RUNX1 (B) and YY1 (C) expression in response to 3 separate shRNA specifically designed to target either RUNX1 or YY1. (D) Principal component analysis was performed on RNA-seq data from AD iNs, AD iNs following RUNX1 KD and AD iNs following YY1 KD (*n* = 3 independent cell lines per group). PC2 and PC3 captured the largest variance among the samples. (E-H) Gating strategy. Example plots and gating for FACS-based purification of the AD iNs and the AD iNs transduced with either SCRM shRNA, RUNX1shRNA or YY1shRNA. iNs were sorted by PSA-NCAM APC and subsequently by mCherry. (I-J) Percentages of FACSorted PSA-NCAM+ cells and mCherry+ PSA-NCAM+ cells over live single cells (*n* = 3 x AD iNs (SCRM), *n* = 3 x AD iNs upon YY1 KD, *n* = 3 x AD iNs upon RUNX1 KD). Statistical significance was assessed by performing one-way ANOVA, Tukey’s multiple comparisons test.

## References

1. Grubman, A. et al. A single-cell atlas of entorhinal cortex from individuals with Alzheimer’s disease reveals cell-type-specific gene expression regulation. Nat Neurosci 22, 2087–2097 (2019).

2. Mathys, H. et al. Single-cell atlas reveals correlates of high cognitive function, dementia, and resilience to Alzheimer’s disease pathology. Cell 186, 4365–4385.e27 (2023).

3. Mertens, J. et al. Age-dependent instability of mature neuronal fate in induced neurons from Alzheimer’s patients. Cell Stem Cell 1533–1548 (2021) doi:10.1016/j.stem.2021.04.004.

4. Renthal, W. et al. Transcriptional Reprogramming of Distinct Peripheral Sensory Neuron Subtypes after Axonal Injury. Neuron 108, 128–144.e9 (2020).

5. Teschendorff, A. E., West, J. & Beck, S. Age-associated epigenetic drift: implications, and a case of epigenetic thrift? Human Molecular Genetics 22, R7–R15 (2013).

6. Traxler, L. et al. Neural cell state shifts and fate loss in ageing and age-related diseases. Nat Rev Neurol 19, 434–443 (2023).

7. Traxler, L. et al. Metabolism navigates neural cell fate in development, aging and neurodegeneration. Dis Model Mech 14, dmm048993 (2021).

8. Xiong, X. et al. Epigenomic dissection of Alzheimer’s disease pinpoints causal variants and reveals epigenome erosion. Cell 186, 4422–4437.e21 (2023).

9. Annese, A. et al. Whole transcriptome profiling of Late-Onset Alzheimer’s Disease patients provides insights into the molecular changes involved in the disease. Sci Rep 8, 4282 (2018).

10. Allen, M. et al. Human whole genome genotype and transcriptome data for Alzheimer’s and other neurodegenerative diseases. Sci Data 3, 160089 (2016).

11. Blalock, E. M., Buechel, H. M., Popovic, J., Geddes, J. W. & Landfield, P. W. Microarray analyses of laser-captured hippocampus reveal distinct gray and white matter signatures associated with incipient Alzheimer’s disease. J Chem Neuroanat 42, 118–126 (2011).

12. Blalock, E. M. et al. Gene microarrays in hippocampal aging: statistical profiling identifies novel processes correlated with cognitive impairment. J Neurosci 23, 3807–3819 (2003).

13. Mills, J. D. et al. RNA-Seq analysis of the parietal cortex in Alzheimer’s disease reveals alternatively spliced isoforms related to lipid metabolism. Neurosci Lett 536, 90–95 (2013).

14. Guo, X.-J., Yang, D. & Zhang, X.-Y. Epigenetics recording varied environment and complex cell events represents the origin of cellular aging. J Zhejiang Univ Sci B 20, 550–562 (2019).

15. Yang, J.-H. et al. Loss of epigenetic information as a cause of mammalian aging. Cell 186, 305–326.e27 (2023).

16. Li, Y. & Tollefsbol, T. O. Age-related epigenetic drift and phenotypic plasticity loss: implications in prevention of age-related human diseases. Epigenomics 8, 1637–1651 (2016).

17. Herdy, J. et al. Chemical modulation of transcriptionally enriched signaling pathways to optimize the conversion of fibroblasts into neurons. eLife 8, 1–21 (2019).

18. Huh, C. J. et al. Maintenance of age in human neurons generated by microRNA-based neuronal conversion of fibroblasts. eLife 5, 1–14 (2016).

19. Kim, Y. et al. Mitochondrial Aging Defects Emerge in Directly Reprogrammed Human Neurons due to Their Metabolic Profile. Cell Reports 23, 2550–2558 (2018).

20. Mertens, J. et al. Directly Reprogrammed Human Neurons Retain Aging-Associated Transcriptomic Signatures and Reveal Age-Related Nucleocytoplasmic Defects. Cell Stem Cell 17, 705–718 (2015).

21. Traxler, L., Edenhofer, F. & Mertens, J. Next-generation disease modeling with direct conversion: a new path to old neurons. FEBS Lett 593, 3316–3337 (2019).

22. Victor, M. B. et al. Striatal neurons directly converted from Huntington’s disease patient fibroblasts recapitulate age-associated disease phenotypes. Nat Neurosci 21, 341–352 (2018).

23. Traxler, L. et al. Warburg-like metabolic transformation underlies neuronal degeneration in sporadic Alzheimer’s disease. Cell Metabolism 34, 1248–1263.e6 (2022).

24. Chen, F. et al. YY1 regulates skeletal muscle regeneration through controlling metabolic reprogramming of satellite cells. EMBO J 38, e99727 (2019).

25. Erwin, J. A., Marchetto, M. C. & Gage, F. H. Mobile DNA elements in the generation of diversity and complexity in the brain. Nat Rev Neurosci 15, 497–506 (2014).

26. Fukui, H., Rünker, A., Fabel, K., Buchholz, F. & Kempermann, G. Transcription factor Runx1 is pro-neurogenic in adult hippocampal precursor cells. PLoS ONE 13, e0190789 (2018).

27. Gordon, S., Akopyan, G., Garban, H. & Bonavida, B. Transcription factor YY1: structure, function, and therapeutic implications in cancer biology. Oncogene 25, 1125–1142 (2006).

28. Logan, T. T., Villapol, S. & Symes, A. J. TGF-β superfamily gene expression and induction of the Runx1 transcription factor in adult neurogenic regions after brain injury. PLoS One 8, e59250 (2013).

29. Okuda, T., Nishimura, M., Nakao, M. & Fujita, Y. RUNX1/AML1: a central player in hematopoiesis. Int J Hematol 74, 252–257 (2001).

30. Sood, R., Kamikubo, Y. & Liu, P. Role of RUNX1 in hematological malignancies. Blood 129, 2070–2082 (2017).

31. Halevy, T., Biancotti, J.-C., Yanuka, O., Golan-Lev, T. & Benvenisty, N. Molecular Characterization of Down Syndrome Embryonic Stem Cells Reveals a Role for RUNX1 in Neural Differentiation. Stem Cell Reports 7, 777–786 (2016).

32. Inoue, K., Shiga, T. & Ito, Y. Runx transcription factors in neuronal development. Neural Dev 3, 20 (2008).

33. Marmigère, F. et al. The Runx1/AML1 transcription factor selectively regulates development and survival of TrkA nociceptive sensory neurons. Nat Neurosci 9, 180–187 (2006).

34. Theriault, F. M. et al. Role for Runx1 in the proliferation and neuronal differentiation of selected progenitor cells in the mammalian nervous system. J Neurosci 25, 2050–2061 (2005).

35. Wang, J. W. & Stifani, S. Roles of Runx Genes in Nervous System Development. Adv Exp Med Biol 962, 103–116 (2017).

36. Duffy, J. B., Kania, M. A. & Gergen, J. P. Expression and function of the Drosophila gene runt in early stages of neural development. Development 113, 1223–1230 (1991).

37. Dormand, E. L. & Brand, A. H. Runt determines cell fates in the Drosophila embryonic CNS. Development 125, 1659–1667 (1998).

38. Stifani, N. et al. Suppression of interneuron programs and maintenance of selected spinal motor neuron fates by the transcription factor AML1/Runx1. Proc Natl Acad Sci U S A 105, 6451–6456 (2008).

39. Eriksson, P. S. et al. Neurogenesis in the adult human hippocampus. Nat Med 4, 1313–1317 (1998).

40. Zurkirchen, L. et al. Yin Yang 1 sustains biosynthetic demands during brain development in a stage-specific manner. Nat Commun 10, 2192 (2019).

41. Wang, Y. et al. Yin Yang 1 promotes the Warburg effect and tumorigenesis via glucose transporter GLUT3. Cancer Science 109, 2423–2434 (2018).

42. Aubry, S. et al. Assembly and Interrogation of Alzheimer’s Disease Genetic Networks Reveal Novel Regulators of Progression. PLoS ONE 10, e0120352 (2015).

43. Buenrostro, J. D., Wu, B., Chang, H. Y. & Greenleaf, W. J. ATAC-seq: A Method for Assaying Chromatin Accessibility Genome-Wide. CP Molecular Biology 109, (2015).

44. Buenrostro, J. D., Giresi, P. G., Zaba, L. C., Chang, H. Y. & Greenleaf, W. J. Transposition of native chromatin for fast and sensitive epigenomic profiling of open chromatin, DNA-binding proteins and nucleosome position. Nature Methods 10, 1213–1218 (2013).

45. Arbab, M., Baars, S. & Geijsen, N. Modeling motor neuron disease: the matter of time. Trends Neurosci 37, 642–652 (2014).

46. Handel, A. E. et al. Assessing similarity to primary tissue and cortical layer identity in induced pluripotent stem cell-derived cortical neurons through single-cell transcriptomics. Hum Mol Genet 25, 989–1000 (2016).

47. Luo, C. et al. Cerebral Organoids Recapitulate Epigenomic Signatures of the Human Fetal Brain. Cell Rep 17, 3369–3384 (2016).

48. Patterson, M. et al. Defining the nature of human pluripotent stem cell progeny. Cell Res 22, 178–193 (2012).

49. Fifre, A. et al. Microtubule-associated protein MAP1A, MAP1B, and MAP2 proteolysis during soluble amyloid beta-peptide-induced neuronal apoptosis. Synergistic involvement of calpain and caspase-3. J Biol Chem 281, 229–240 (2006).

50. Gutiérrez-Vargas, J. A., Castro-Álvarez, J. F., Zapata-Berruecos, J. F., Abdul-Rahim, K. & Arteaga-Noriega, A. Neurodegeneration and convergent factors contributing to the deterioration of the cytoskeleton in Alzheimer’s disease, cerebral ischemia and multiple sclerosis (Review). Biomed Rep 16, 27 (2022).

51. Li, B. et al. Failure of neuronal maturation in Alzheimer disease dentate gyrus. J Neuropathol Exp Neurol 67, 78–84 (2008).

52. Vontell, R. T. et al. Association of region-specific hippocampal reduction of neurogranin with inflammasome proteins in post mortem brains of Alzheimer’s disease. Alzheimers Dement (N Y) 10, e12444 (2024).

53. Kim, H. J. et al. REST Regulates Non-Cell-Autonomous Neuronal Differentiation and Maturation of Neural Progenitor Cells via Secretogranin II. J Neurosci 35, 14872–14884 (2015).

54. Joly-Amado, A., Kulkarni, N. & Nash, K. R. Reelin Signaling in Neurodevelopmental Disorders and Neurodegenerative Diseases. Brain Sci 13, 1479 (2023).

55. Lane-Donovan, C. et al. Reelin protects against amyloid β toxicity in vivo. Sci Signal 8, ra67 (2015).

56. Lleó, A. et al. VAMP-2 is a surrogate cerebrospinal fluid marker of Alzheimer-related cognitive impairment in adults with Down syndrome. Alzheimers Res Ther 13, 119 (2021).

57. Roussarie, J.-P. et al. Selective Neuronal Vulnerability in Alzheimer’s Disease: A Network-Based Analysis. Neuron 107, 821–835.e12 (2020).

58. Taddei, R. N. & E Duff, K. Synapse vulnerability and resilience underlying Alzheimer’s disease. EBioMedicine 112, 105557 (2025).

59. Williams, J. B., Cao, Q. & Yan, Z. Transcriptomic analysis of human brains with Alzheimer’s disease reveals the altered expression of synaptic genes linked to cognitive deficits. Brain Commun 3, fcab123 (2021).

60. Furlan, F. et al. Interneurons transiently express the ERG K+ channels during development of mouse spinal networks in vitro. Neuroscience 135, 1179–1192 (2005).

61. Kim, D.-Y., Hwang, I., Muller, F. L. & Paik, J.-H. Functional regulation of FoxO1 in neural stem cell differentiation. Cell Death Differ 22, 2034–2045 (2015).

62. Komorowska, K. et al. Hepatic Leukemia Factor Maintains Quiescence of Hematopoietic Stem Cells and Protects the Stem Cell Pool during Regeneration. Cell Rep 21, 3514–3523 (2017).

63. Lehnertz, B. et al. HLF expression defines the human hematopoietic stem cell state. Blood 138, 2642–2654 (2021).

64. Liu, L. et al. Cross-Talking Pathways of Forkhead Box O1 (FOXO1) Are Involved in the Pathogenesis of Alzheimer’s Disease and Huntington’s Disease. Oxid Med Cell Longev 2022, 7619255 (2022).

65. Loughran, S. J. et al. The transcription factor Erg is essential for definitive hematopoiesis and the function of adult hematopoietic stem cells. Nat Immunol 9, 810–819 (2008).

66. Lu, H. & Huang, H. FOXO1: a potential target for human diseases. Curr Drug Targets 12, 1235–1244 (2011).

67. Angel, P. & Karin, M. The role of Jun, Fos and the AP-1 complex in cell-proliferation and transformation. Biochim Biophys Acta 1072, 129–157 (1991).

68. Karin, M., Liu, Z. g & Zandi, E. AP-1 function and regulation. Curr Opin Cell Biol 9, 240–246 (1997).

69. Martínez-Zamudio, R. I. et al. AP-1 imprints a reversible transcriptional programme of senescent cells. Nat Cell Biol 22, 842–855 (2020).

70. Zumerle, S. & Alimonti, A. In and out from senescence. Nat Cell Biol 22, 753–754 (2020).

71. Marciniak, S. J. et al. CHOP induces death by promoting protein synthesis and oxidation in the stressed endoplasmic reticulum. Genes Dev 18, 3066–3077 (2004).

72. Wortel, I. M. N., van der Meer, L. T., Kilberg, M. S. & van Leeuwen, F. N. Surviving Stress: Modulation of ATF4-Mediated Stress Responses in Normal and Malignant Cells. Trends Endocrinol Metab 28, 794–806 (2017).

73. Wu, D. & Liang, J. Activating transcription factor 4: a regulator of stress response in human cancers. Front Cell Dev Biol 12, 1370012 (2024).

74. Zinszner, H. et al. CHOP is implicated in programmed cell death in response to impaired function of the endoplasmic reticulum. Genes Dev 12, 982–995 (1998).

75. Chatterjee, N. & Walker, G. C. Mechanisms of DNA damage, repair, and mutagenesis. Environ Mol Mutagen 58, 235–263 (2017).

76. Citron, B. A., Dennis, J. S., Zeitlin, R. S. & Echeverria, V. Transcription factor Sp1 dysregulation in Alzheimer’s disease. J Neurosci Res 86, 2499–2504 (2008).

77. Wang, L. et al. The effects of nitric oxide in Alzheimer’s disease. Med Gas Res 14, 186–191 (2024).

78. Hänzelmann, S. et al. Replicative senescence is associated with nuclear reorganization and with DNA methylation at specific transcription factor binding sites. Clin Epigenetics 7, 19 (2015).

79. Herdy, J. R. et al. Increased post-mitotic senescence in aged human neurons is a pathological feature of Alzheimer’s disease. Cell Stem Cell 29, 1637–1652.e6 (2022).

80. Ohtani, N. et al. Opposing effects of Ets and Id proteins on p16INK4a expression during cellular senescence. Nature 409, 1067–1070 (2001).

81. Anderson, S. R. et al. Disrupted SOX10 function causes spongiform neurodegeneration in gray tremor mice. Mamm Genome 26, 80–93 (2015).

82. Casalino, L., Talotta, F., Cimmino, A. & Verde, P. The Fra-1/AP-1 Oncoprotein: From the ‘Undruggable’ Transcription Factor to Therapeutic Targeting. Cancers (Basel) 14, 1480 (2022).

83. Fang, X. et al. Twist2 contributes to breast cancer progression by promoting an epithelial-mesenchymal transition and cancer stem-like cell self-renewal. Oncogene 30, 4707–4720 (2011).

84. Foletta, V. C. Transcription factor AP-1, and the role of Fra-2. Immunol Cell Biol 74, 121–133 (1996).

85. Jeter, C. R., Yang, T., Wang, J., Chao, H.-P. & Tang, D. G. Concise Review: NANOG in Cancer Stem Cells and Tumor Development: An Update and Outstanding Questions. Stem Cells 33, 2381–2390 (2015).

86. Leiherer, A., Geiger, K., Muendlein, A. & Drexel, H. Hypoxia induces a HIF-1α dependent signaling cascade to make a complex metabolic switch in SGBS-adipocytes. Mol Cell Endocrinol 383, 21–31 (2014).

87. Lorenzin, F. & Demichelis, F. Past, Current, and Future Strategies to Target ERG Fusion-Positive Prostate Cancer. Cancers (Basel) 14, 1118 (2022).

88. Pérez-Benavente, B. et al. New roles for AP-1/JUNB in cell cycle control and tumorigenic cell invasion via regulation of cyclin E1 and TGF-β2. Genome Biol 23, 252 (2022).

89. Renoux, F. et al. The AP1 Transcription Factor Fosl2 Promotes Systemic Autoimmunity and Inflammation by Repressing Treg Development. Cell Rep 31, 107826 (2020).

90. Thompson, M. R., Xu, D. & Williams, B. R. G. ATF3 transcription factor and its emerging roles in immunity and cancer. J Mol Med (Berl) 87, 1053–1060 (2009).

91. Duclot, F. & Kabbaj, M. The Role of Early Growth Response 1 (EGR1) in Brain Plasticity and Neuropsychiatric Disorders. Front Behav Neurosci 11, 35 (2017).

92. Knapska, E. & Kaczmarek, L. A gene for neuronal plasticity in the mammalian brain: Zif268/Egr-1/NGFI-A/Krox-24/TIS8/ZENK? Prog Neurobiol 74, 183–211 (2004).

93. Qin, X., Wang, Y. & Paudel, H. K. Inhibition of Early Growth Response 1 in the Hippocampus Alleviates Neuropathology and Improves Cognition in an Alzheimer Model with Plaques and Tangles. Am J Pathol 187, 1828–1847 (2017).

94. Heinrich, R., Aronheim, A., Cheng, Y.-C. & Perlman, I. Editorial: ATF3: a crucial stress-responsive gene of glia and neurons in CNS. Front Mol Neurosci 17, 1484487 (2024).

95. Lu, W. et al. Over-expression of c-fos mRNA in the hippocampal neurons in Alzheimer’s disease. Chin Med J (Engl) 111, 35–37 (1998).

96. Soni, P., Sharma, S. M., Pieper, A. A., Paul, B. D. & Thomas, B. Nrf2/Bach1 signaling axis: A promising multifaceted therapeutic strategy for Alzheimer’s disease. Neurotherapeutics 22, e00586 (2025).

97. Wei, X. et al. The Multifaceted Roles of BACH1 in Disease: Implications for Biological Functions and Therapeutic Applications. Adv Sci (Weinh) 12, e2412850 (2025).

98. Yang, T. et al. The potential roles of ATF family in the treatment of Alzheimer’s disease. Biomed Pharmacother 161, 114544 (2023).

99. Bai, B. et al. Deep Multilayer Brain Proteomics Identifies Molecular Networks in Alzheimer’s Disease Progression. Neuron 105, 975–991.e7 (2020).

100. Johnson, E. C. B. et al. Large-scale proteomic analysis of Alzheimer’s disease brain and cerebrospinal fluid reveals early changes in energy metabolism associated with microglia and astrocyte activation. Nat Med 26, 769–780 (2020).

101. Cao, Y. Tumorigenesis as a process of gradual loss of original cell identity and gain of properties of neural precursor/progenitor cells. Cell Biosci 7, 61 (2017).

102. Corces, M. R. et al. Lineage-specific and single-cell chromatin accessibility charts human hematopoiesis and leukemia evolution. Nat Genet 48, 1193–1203 (2016).

103. Friedmann-Morvinski, D. & Verma, I. M. Dedifferentiation and reprogramming: origins of cancer stem cells. EMBO Rep 15, 244–253 (2014).

104. Poplawski, G. H. D. et al. Injured adult neurons regress to an embryonic transcriptional growth state. Nature 581, 77–82 (2020).

105. Yzaguirre, A. D., de Bruijn, M. F. T. R. & Speck, N. A. The Role of Runx1 in Embryonic Blood Cell Formation. Adv Exp Med Biol 962, 47–64 (2017).

106. Burns, C. E., Traver, D., Mayhall, E., Shepard, J. L. & Zon, L. I. Hematopoietic stem cell fate is established by the Notch-Runx pathway. Genes Dev 19, 2331–2342 (2005).

107. Ito, Y. & Miyazono, K. RUNX transcription factors as key targets of TGF-beta superfamily signaling. Curr Opin Genet Dev 13, 43–47 (2003).

108. Asou, N. The role of a Runt domain transcription factor AML1/RUNX1 in leukemogenesis and its clinical implications. Crit Rev Oncol Hematol 45, 129–150 (2003).

109. Ito, Y. RUNX genes in development and cancer: regulation of viral gene expression and the discovery of RUNX family genes. Adv Cancer Res 99, 33–76 (2008).

110. Miyoshi, H. et al. t(8;21) breakpoints on chromosome 21 in acute myeloid leukemia are clustered within a limited region of a single gene, AML1. Proc Natl Acad Sci U S A 88, 10431–10434 (1991).

111. De Vita, S. et al. Trisomic dose of several chromosome 21 genes perturbs haematopoietic stem and progenitor cell differentiation in Down’s syndrome. Oncogene 29, 6102–6114 (2010).

112. Liu, Y., Zhang, Y., Ren, Z., Zeng, F. & Yan, J. RUNX1 Upregulation Causes Mitochondrial Dysfunction via Regulating the PI3K-Akt Pathway in iPSC from Patients with Down Syndrome. Mol Cells 46, 219–230 (2023).

113. Raha-Chowdhury, R. et al. Impaired Iron Homeostasis and Haematopoiesis Impacts Inflammation in the Ageing Process in Down Syndrome Dementia. J Clin Med 10, 2909 (2021).

114. Vilardell, M. et al. Meta-analysis of heterogeneous Down Syndrome data reveals consistent genome-wide dosage effects related to neurological processes. BMC Genomics 12, 229 (2011).

115. Patel, A. et al. Association of variants within APOE, SORL1, RUNX1, BACE1 and ALDH18A1 with dementia in Alzheimer’s disease in subjects with Down syndrome. Neuroscience Letters 487, 144–148 (2011).

116. He, Y. et al. The transcription factor Yin Yang 1 is essential for oligodendrocyte progenitor differentiation. Neuron 55, 217–230 (2007).

117. He, Y. & Casaccia-Bonnefil, P. The Yin and Yang of YY1 in the nervous system. J Neurochem 106, 1493–1502 (2008).

118. Liu, H. et al. Yin Yang 1 is a critical regulator of B-cell development. Genes Dev 21, 1179–1189 (2007).

119. Verheul, T. C. J., van Hijfte, L., Perenthaler, E. & Barakat, T. S. The Why of YY1: Mechanisms of Transcriptional Regulation by Yin Yang 1. Front Cell Dev Biol 8, 592164 (2020).

120. Weintraub, A. S. et al. YY1 Is a Structural Regulator of Enhancer-Promoter Loops. Cell 171, 1573–1588.e28 (2017).

121. Chen, Z. S. & Chan, H. Y. E. Transcriptional dysregulation in neurodegenerative diseases: Who tipped the balance of Yin Yang 1 in the brain? Neural Regen Res 14, 1148–1151 (2019).

122. Garcia-Alonso, L., Holland, C. H., Ibrahim, M. M., Turei, D. & Saez-Rodriguez, J. Benchmark and integration of resources for the estimation of human transcription factor activities. Genome Res 29, 1363–1375 (2019).

123. Heinz, S. et al. Simple combinations of lineage-determining transcription factors prime cis-regulatory elements required for macrophage and B cell identities. Mol Cell 38, 576–589 (2010).

124. Ramírez, F. et al. deepTools2: a next generation web server for deep-sequencing data analysis. Nucleic Acids Res 44, W160–165 (2016).

